# Mechanism of *ATXN8OS* CTA/CTG repeat-associated non-AUG translation revealed by approaches ranging from cell-free translation to live-cell imaging

**DOI:** 10.64898/2026.01.25.700793

**Authors:** Soyoka Sakamoto, Hayato Ito, Mayuka Hasumi, Tatsuya Morisaki, Makito Hirano, Yoshitaka Nagai, Timothy J. Stasevich, Hideki Taguchi

## Abstract

Microsatellite repeat expansions contribute to the pathogenesis of many neurodegenerative disorders. In spinocerebellar ataxia type 8 (SCA8), abnormal expansion of CTA/CTG repeats in the 3′ untranslated region of the *ATXN8OS* (*ATXN8* Opposite Strand) gene has been implicated in disease pathology. Although the occurrence of repeat-associated non-AUG (RAN) translation from the *ATXN8* transcript has been reported, whether and how RAN translation occurs from the *ATXN8OS* transcript has remained unexplored. Here, using a cell-free translation system and cultured cells, we showed that *ATXN8OS* undergoes robust AUG-independent translation in a repeat length–dependent manner. Mechanistic analyses revealed that translation of the poly L (0) frame initiates at a non-AUG codon located upstream of the repeats. Moreover, using live-cell imaging at a single mRNA level, we directly visualized ribosomal −1 frameshifting from the poly L (0) frame to the poly T-poly A (+2) frame during translation elongation. We further showed that *ATXN8OS* translation was enhanced upon activation of the integrated stress response. Together, these findings establish both the occurrence and the molecular mechanisms of *ATXN8OS* translation from expanded CTA/CTG repeats and provide insights into the pathogenic processes underlying SCA8.

**GRAPHICAL ABSTRACT:** 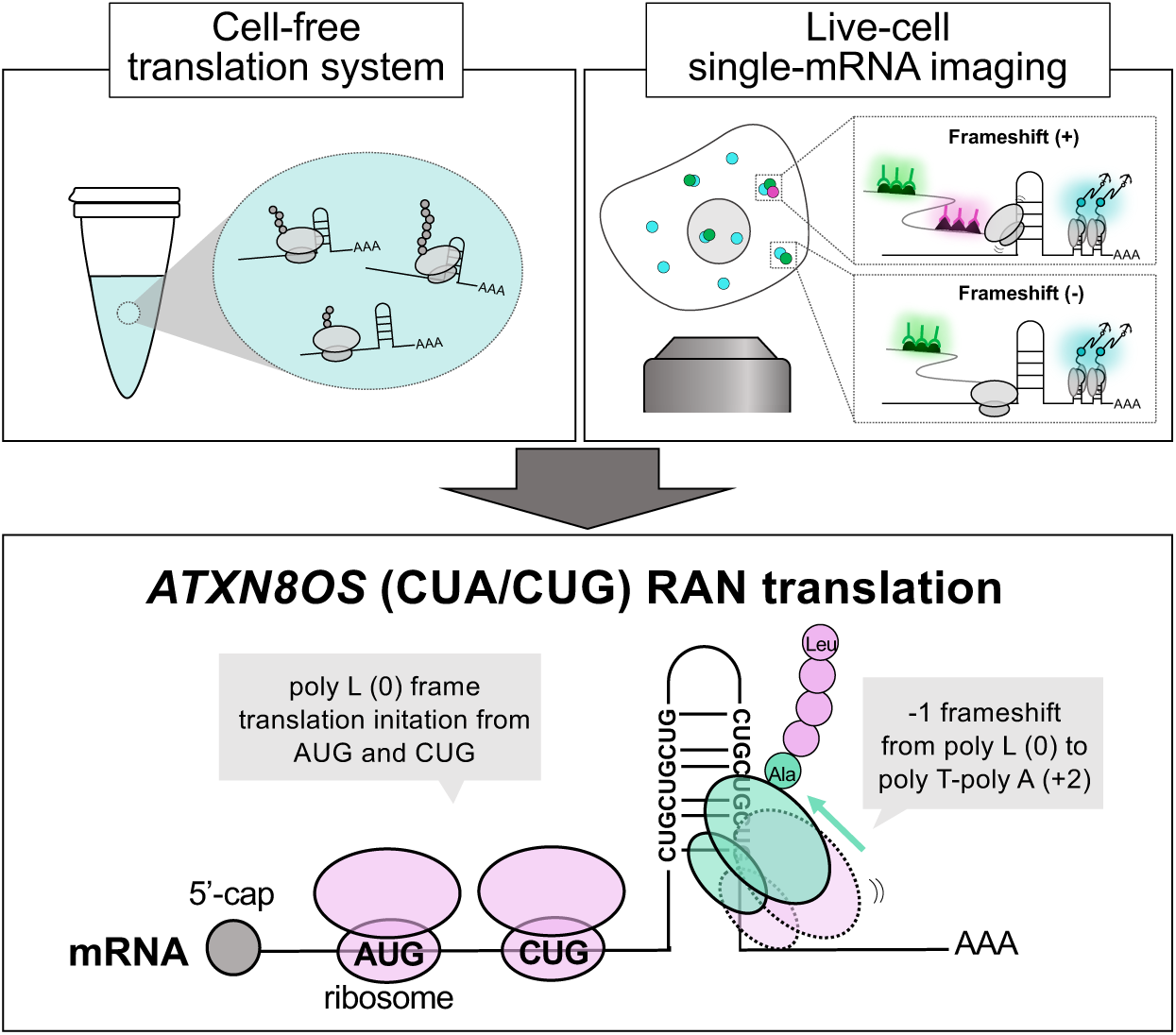

## Introduction

Short tandem repeat expansions are a major genetic cause of inherited neurodegenerative diseases (1,2). Repeat expansions were initially thought to drive diseases primarily through two mechanisms: loss of corresponding gene function and RNA-mediated toxicity (3). More recently, however, studies have revealed that a non-canonical translation mechanism, termed repeat-associated non-AUG (RAN) translation, is related to disease pathogenesis (4–7). RAN translation occurs on abnormal transcripts harboring expanded repeats, which are translated in multiple reading frames in the absence of a canonical AUG start codon, resulting in the production of toxic repeat peptides (8). RAN translation has been identified across numerous repeat expansion disorders and has therefore emerged as a potential therapeutic target (9).

At the mechanistic level, RAN translation is driven by structured, GC-rich RNA sequences, such as G-quadruplexes or stable hairpins, and proceeds without a canonical AUG initiation codon (10). A defining feature of RAN translation is its ability to initiate in all three reading frames. Since its initial discovery in spinocerebellar ataxia type 8 (SCA8) and myotonic dystrophy type 1 (DM1) (4), RAN translation has been reported in a wide range of repeat expansion disorders (5–7,11–14), although the underlying molecular mechanisms remain incompletely understood. With respect to translation initiation, several studies have shown that RAN translation can occur via cap-dependent scanning mediated by eIF4E and eIF4A, initiating from near-cognate codons located upstream of the repeat, as exemplified by *C9orf72* (GGGGCC; C9-RAN), *FMR1* (CGG; FMR1-RAN), and *Nop56* (GGCCTG; NOP56-RAN) (15–17). In other cases, non-cognate non-AUG codons are used; for example, *ATXN3* (CAG; SCA3-RAN) utilizes CUU and ACU codons (11). In addition, C9-RAN has also been proposed to occur in a 5′ cap-independent manner, resembling an internal ribosome entry site (IRES) (18–20). Importantly, these non-canonical initiation events are strongly enhanced by increasing repeat length (4,16,17,21). Beyond initiation, RAN translation is also regulated at the elongation stage. Longer repeat RNAs stimulate ribosomal frameshifting (17,21–24). The frameshifting is thought to be promoted by repetitive codons and stable RNA structures (25,26). Moreover, GC-rich RNA repeats stall elongating ribosomes and activate ribosome-associated quality control pathways (27). Finally, RAN translation is upregulated upon activation of the integrated stress response (ISR), such as ER stress via the PERK pathway (11,15,17,28), providing a mechanistic link between cellular stress and repeat-associated disease pathogenesis.

SCA8 is associated with an abnormal expansion of CTA/CTG repeats in the 3′ untranslated region (UTR) of the *ATXN8OS* (*ATXN8* Opposite Strand) gene (Fig. 1, Supplementary Fig. S1A) (29). SCA8 is a slowly progressive, adult-onset cerebellar ataxia characterized by marked cerebellar atrophy, manifesting as gait and speech disturbances and, in some cases, parkinsonism (30–32). The normal CTA/CTG repeat length ranges from 15 to 50 repeats (rp), whereas expansions of 71 to 1300 repeats are considered pathogenic (3,29). Notably, a recent study further reported that approximately 3% of Japanese amyotrophic lateral sclerosis (ALS) patients carry a CTA/CTG repeat expansion in *ATXN8OS* (33).

**Figure 1.**
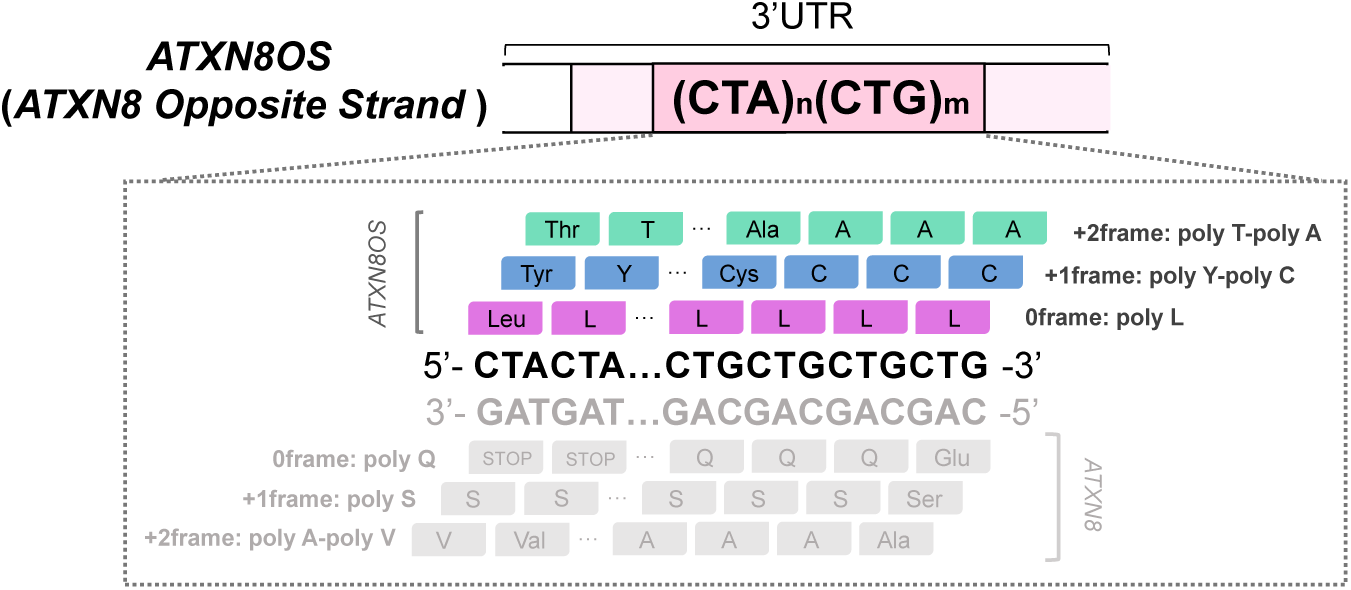
Putative repeat peptides encoded by the *ATXN8OS* CUA/CUG repeat. Schematic of putative translation products derived from the *ATXN8OS* CUA/CUG repeat. *ATXN8OS* is predicted to encode poly L (0 frame), poly Y-poly C (+1 frame), and poly T-poly A (+2 frame) peptides.

The CTA/CTG repeat expansion is transcribed from both strands, producing polyadenylated *ATXN8OS*-derived CUA/CUG repeat RNA (*ATXN8OS* RNA) and *ATXN8*-derived CAG/UAG containing RNA (*ATXN8* RNA) (34,35). Previous studies have shown that *ATXN8OS* RNA mediates RNA toxicity by sequestering RNA-binding proteins, including the splicing factor MBNL1 (36,37). In parallel, *ATXN8* RNA encodes an ATG-initiated polyglutamine (poly Q) protein that also contributes to disease pathogenesis (35).

In SCA8, studies using the *ATXN8* CAG repeat demonstrated AUG-independent translation in all three reading frames (0 frame: poly Q, +1 frame: poly S, +2 frame: poly A), which led to the proposal of RAN translation (4). In contrast to the extensive characterization of *ATXN8* RNA, research on *ATXN8OS* RNA has remained limited, despite clear evidence of its transcription. Indeed, several studies failed to detect appreciable levels of translation products from *ATXN8OS* (29,35). Even in cases where RAN translation was detected, analyses were restricted to the CUG repeat region within the *ATXN8OS* sequence (4).

To elucidate the molecular mechanisms underlying *ATXN8OS*-derived CUA/CUG translation in its native sequence context (Supplementary Fig. S1B), we combined multiple experimental approaches. Building on our recent work, we analyzed translation using both cell-free and cell-based reporter assays (17,21). We detected robust RAN translation from *ATXN8OS*. In addition, we employed live-cell imaging to directly visualize ribosomal frameshifting events at a single mRNA level (22,38,39). Together, these experiments define key features of *ATXN8OS* translation, including its initiation sites, repeat length dependence, ribosomal frameshifting, and response to ER stress. Collectively, these findings provide new mechanistic insight into *ATXN8OS* translation and establish a framework for understanding its contribution to SCA8 pathogenesis.

## Materials and Methods

### Plasmids

All plasmids were generated using standard cloning procedures followed by Gibson assembly. To maintain the stability of repeat sequences, all constructs were cloned in SURE2 cells. For example, the *ATXN8OS*– nano-luciferase (Nluc) poly L (0) frame reporter plasmid for in vitro translation was constructed by PCR amplifying the 3′ UTR region of *ATXN8OS* and the pcDNA5/T7-NOP56-Nluc+3×HA vector backbone (17). The *ATXN8OS* 3′ UTR was amplified using primers SS007 (GAATTGCCCTTTAAGTAACTAAACGGAGAGATTAACTCTGTTGGCTGAAGCCCTATT) and SS009 (ATCTTCGAGTGTGAAGACCCCAAATAGTAAAATAAGATAATATATTTGTAAAAAATGCAGCAGCA), and the vector was amplified using primers SS002 (GTCTTCACACTCGAAGATTTCGTTGG) and SS001 (TTAGTTACTTAAAGGGCAATTCCACCACACTGGAC). PCR amplification was performed using PrimeSTAR Max DNA Polymerase (Takara). For constructs containing the 80-repeat (80 rp) insert, betaine was added at 32.5% (v/v) to stabilize the repeat. PCR conditions were as follows: 35 cycles of 10 s at 95 °C, 5 s at 58 °C, and 25 s at 72 °C, followed by a final extension for 5 min at 72 °C. PCR products were verified by agarose gel electrophoresis for correct size. Insert and vector fragments were subsequently assembled using Gibson assembly.

Sanger sequencing was performed to verify sequence accuracy. *ATXN8OS* CUA/CUG repeats of various lengths were obtained by PCR. Reporter plasmids for cell-based assays were generated by inserting the 3′ UTR region of *ATXN8OS* into the pcDNA-FRT-T7-C9ORF72-Nluc-3×FLAG vector (40). Reporter plasmids for immunostaining were generated by inserting the 3′ UTR region of *ATXN8OS* into the pcDNA5/FRT-T7-3FT vector (21). Initiation codon-mutated reporters and Kozak sequence-mutated reporters were generated using PCR and Gibson assembly. For the frameshift-detection reporter, an ATG–FLAG sequence was introduced into pcDNA-FRT-T7-ATXN8OS-Nluc-3×HA. For live-cell imaging reporters, a spaghetti monster HA tag (smHA) was first inserted upstream of the *ATXN8OS* repeat sequence in the poly L (0) frame. The smHA plasmid (Addgene plasmid #81085) and the MoonTag and SunTag hybrid (MASH) reporter cassette (Addgene plasmid #128607) were inserted downstream of the *ATXN8OS* repeat sequence (41). In this construct, the smHA was placed in the poly L (0) frame, whereas the SunTag was positioned in the poly T–poly A (+2) frame. All reporter constructs were generated using Gibson assembly. To enable nascent chain tracking, an all-probes plasmid was generated by modifying the indicated backbone plasmid (Addgene plasmid #241139) (42). Briefly, the fluorescent protein mBaoJin (43), synthesized by Twist Bioscience, was inserted downstream of the HA-Tag Frankenbody (44) (Addgene plasmid #129591), and mScarlet was inserted downstream of the anti-SunTag (GCN4) single-chain variable fragment (scFv). In addition, to visualize mRNA molecules, the PP7 coat protein gene was inserted upstream of a HaloTag protein containing a C-terminal CaaX motif for plasma membrane tethering. This design allowed simultaneous visualization of HA- and SunTag-labeled nascent polypeptides together with PP7-tagged mRNA. All plasmid sequences were verified by Sanger sequencing. Supplementary Tables S1 and S2 list all plasmids and oligonucleotides, respectively, used in this study.

### In vitro transcription

In vitro transcription was carried out following previously established protocols (17,21). Reporter plasmids were linearized using XbaI (Takara). The resulting fragments were purified with the Wizard SV Gel and PCR Purification Kit (Promega). 5′-capped mRNAs were generated using the mMESSAGE mMACHINE T7 Transcription Kit (Thermo Fisher Scientific), and then polyadenylated with *Escherichia coli* poly(A) polymerase (NEB). Uncapped mRNAs were generated using the CUGA T7 in vitro transcription kit (Nippon Gene). All mRNAs were purified by lithium chloride precipitation. Their size was assessed by denaturing agarose gel electrophoresis, and concentrations were measured by NanoDrop spectrophotometry (Thermo Fisher Scientific).

### In vitro translation in HeLa lysate

In vitro translation was carried out using HeLa cell lysates as previously described (17,21,45). Reactions were assembled at a final mRNA concentration of 10 nM, incubated at 32 °C for 1.5 h, and then halted by placing the samples on ice. For free m⁷GpppG compound addition assays, the cap analog was supplemented at a concentration of 2.5 mM. After the reaction, 2 µL of each sample was diluted in Glo Lysis Buffer (Promega) in a 96-well plate. The diluted samples were then mixed at a 1:1 ratio with freshly prepared NanoGlo substrate (1:50 dilution in NanoGlo buffer; Promega) and incubated for 3 min with shaking in the dark. Luminescence signals were recorded using a Varioskan LUX Multimode Microplate Reader (Thermo Fisher Scientific). For western blotting, the same translation reactions were processed, and 6 µL of each sample was used. Samples were mixed with 15 µL of 4× SDS sample buffer (240 mM Tris–HCl, pH 6.8; 40% (v/v) glycerol; 0.01% (w/v) bromophenol blue; 7% (w/v) SDS; 10% (v/v) 2-mercaptoethanol) and heated at 70 °C for 15 min. Subsequently, 15 µL of each denatured sample was loaded onto a 13% SDS–polyacrylamide gel for electrophoresis, followed by western blotting.

### In vitro translation in rabbit reticulocyte lysate (RRL)

In vitro translation using rabbit reticulocyte lysate (RRL) was carried out as previously described (17,46). Reactions were assembled at a final mRNA concentration of 3 nM, incubated at 30 °C for 30 min, and then halted by placing the samples on ice. The resulting samples were processed using the same workflow described for in vitro translation in HeLa lysate.

### Cell culture and transfections

HeLa cells, HEK293T cells, and U2OS cells were all cultured in Dulbecco’s Modified Eagle Medium (DMEM; Nacalai Tesque), supplemented with 2 mM L-glutamine, 100 U/mL penicillin, 0.1 mg/mL streptomycin (Sigma-Aldrich), and 10% (v/v) fetal bovine serum (FBS; Thermo Fisher Scientific). For the dual-luciferase assays, cells were seeded at 1.0 × 10^4^ cell per well in 96-well plates.

After incubating for 24 h, cells were transfected with 50 ng of a Fluc control plasmid and 50 ng of an *ATXN8OS*-Nluc reporter plasmid using 0.3 µL FuGENE HD (Promega) diluted in 20 µL Opti-MEM (Invitrogen) per well. 24 h after transfection, cells were lysed in 50 µL Glo Lysis Buffer (Promega). For stress induction, HEK293T cells were transfected for 19 h and then treated with 1 µM thapsigargin (Thermo Fisher Scientific, T7458) for 5 h before lysis. After lysis, 30 µL of lysate was mixed 1:1 with ONE-Glo EX substrate and incubated for 3 min with shaking in the dark. Luminescence was quantified using a Varioskan LUX Multimode Microplate Reader (Thermo Fisher Scientific). After Fluc detection, 30 µL of NanoGlo substrate (1:100 dilution in NanoGlo buffer; Promega) was added to the same wells, shaken for 3 min in the dark, and Nluc activity was measured. For western blotting, cells were seeded at 6 × 10^5^ cells per well in 12-well plates. After incubating for 24 h, cells were transfected with 500 ng of *ATXN8OS*-Nluc reporter plasmid using 1.4 µL FuGENE HD and 50 µL Opti-MEM per well. 24 h after transfection, cells were lysed in 150 µL RIPA buffer (50 mM Tris-HCl pH 6.8, 150 mM NaCl, 0.1% (w/v) SDC, 0.1% (w/v) SDS, 1.0% (w/v) NP-40). 70 µL of the lysates was mixed with 70 µL 4 × SDS sample buffer and heated at 70 °C for 15 min. Subsequently, 15 µL of each denatured sample was loaded onto a 13% SDS–polyacrylamide gel for electrophoresis, followed by Western blotting. For immunostaining, cells were seeded at 1 × 10^4^ cells per well in 24-well glass bottom plates. After incubating for 24 h, cells were transfected with 250 ng of *ATXN8OS* reporter plasmid using 1.5 µL FuGENE HD and 100 µL Opti-MEM per well. Cells were incubated for 48 h prior to fixation.

### Western Blotting

Protein samples were separated by size on 13% SDS–polyacrylamide gels and subsequently transferred to PVDF membranes. Membranes were blocked in 2% (w/v) skim milk in TBS-T for 1 h and then incubated with the primary antibodies listed in Supplementary Table S3 for 1 h. After washing with TBS-T, membranes were incubated with an HRP-conjugated anti-mouse IgG secondary antibody (Sigma-Aldrich) for 1 h Chemiluminescent signals were visualized using an LAS4000 imaging system (Fujifilm).

### Immunostaining

48 h after transfection, HeLa cells were fixed with 4% paraformaldehyde in 250 mM HEPES-KOH pH 7.5 for 20 min. Subsequently, cells were permeabilized by shaking for 20 min with 1% Triton X-100 in PBS. After blocking with Blocking One-P (Nacalai Tesque) for 20 min, cells were incubated for 2 h with the primary antibodies listed in Supplementary Table S3. After washing with PBS, cells were incubated for 1.5 h with the secondary antibodies listed in Supplementary Table S3. Imaging was performed using a spinning-disk confocal microscope (Dragonfly; Andor). The Dragonfly system is equipped with a laser unit comprising 405, 488, 561, and 637 nm excitation lines and is mounted on an inverted microscope (Ti-E; Nikon). Microscope control and image acquisition were carried out using Fusion software (version 2.3.0.45; Andor). Images were acquired using a 100× oil-immersion objective lens (Plan Apo VC, NA 1.4; Nikon).

### Live-cell imaging

U2OS cells were seeded at 1.0 × 10^4^ cells per 35-mm glass-bottom dish and cultured in Opti-MEM (Invitrogen) supplemented with 2 mM L-glutamine, 100 U/mL penicillin, 0.1 mg/mL streptomycin (Sigma-Aldrich), and 10% (v/v) FBS (Thermo Fisher Scientific). After incubating for 24 h, cells were transfected with 300 ng of *ATXN8OS* reporter plasmid and 500 ng of all-probes plasmid using 0.9 µL PLUS reagent and 2.7 µL Lipofectamine LTX (Thermo Fisher Scientific), diluted in 50 µL Opti-MEM (Invitrogen) per dish. 6 h after transfection, the culture medium was replaced with FluoroBrite DMEM (Thermo Fisher Scientific) supplemented with 2 mM L-glutamine, 100 U/mL penicillin, 0.1 mg/mL streptomycin (Sigma-Aldrich), and 10% (v/v) FBS (Thermo Fisher Scientific), and cells were incubated with 100 nM Janelia Fluor Halo dye (JF646) (Promega). After 30 min incubation, JF646 was washed out, and cells were incubated for >1 h with FluoroBrite DMEM prior to imaging. Images were acquired on a custom-built HILO microscope (39,47). Briefly, the excitation beams (488, 561, and 637 nm solid-state lasers; Vortran) were coupled and focused at the back focal plane of the objective lens (60×, NA 1.49 oil-immersion objective, Olympus). The emission signals were split by an imaging-grade, ultra-flat dichroic mirror (T660lpxr, Chroma) and detected using an EM-CCD camera (iXon Ultra 888; Andor) via focusing with 300-mm tube lenses (producing 100× images with 130 nm/pixel). Live cells were maintained in a stage-top incubator at 37 °C with 5% CO₂ (Okolab) and imaged on a piezoelectric stage (PZU-2150; Applied Scientific Instrumentation). Focus was maintained throughout image acquisition using the CRISP autofocus system (CRISP-890; Applied Scientific Instrumentation). Laser emission, camera integration, piezoelectric stage movement, and emission filter wheel switching were synchronized using an Arduino Mega controller (Arduino). Image acquisition and hardware control (lasers, cameras, filter wheel, and piezo stage) were performed using pycro-manager, a Python-based interface to Micro-Manager (48). Images were acquired with a 512 × 512 pixel region of interest, corresponding to an imaging area of 66.6 × 66.6 µm. The exposure time was set to 53.64 ms. The EM-CCD camera readout time was 23.36 ms, resulting in an effective imaging rate of approximately 13 Hz (∼70 ms per frame). Excitation lasers were digitally synchronized with camera exposure to minimize photobleaching. To capture the full cytoplasmic volume of U2OS cells, 13 z-planes were acquired with a step size of 500 nm, corresponding to a total z-range of 6 µm.

### Tracking translation of single mRNAs

Image pre-processing was performed using Fiji by generating maximum-intensity Z-projections for each time frame of the three-dimensional movies. Single mRNA particles were tracked using the TrackMate plugin (49). The resulting data were subsequently analyzed using custom-written Python code to assess translation status in a semi-automated manner. Specifically, a maximum-projection was applied to the mRNA channel across the Z-stack, and mRNA particles were then tracked using the TrackMate plugin (50). Detection was performed with the LoG (Laplacian of Gaussian) detector, with the particle diameter set to 7 pixels. Detection thresholds were manually adjusted for each image. Detected particles were tracked using the simple LAP (Linear Assignment Problem) tracker, with a maximum linking distance of 3 pixels, a gap-closing distance of 5 pixels, and a maximum gap of 2 frames. To reduce noise-driven trajectories, only tracks persisting for at least 12 frames (1 min) were included in subsequent analysis. To calculate translation-site intensities, the acquired images were first corrected for photobleaching according to previously published procedures (42). For each detected translation site, crops were generated based on the spot coordinates obtained from the TrackMate output tracking CSV files. Crops from each track were manually inspected to ensure high signal quality with clearly visible spots in all frames. Translation intensities were then quantified using the Disk–Donut method (38,42). Briefly, the signal intensity was calculated as the average pixel intensity within a centered disk of 3-pixel radius, and local background was estimated from a surrounding annular region with an inner radius of 5 pixels and a width of 1 pixel. The background-subtracted intensity was used as the translation-site signal. Subsequently, spot quality was manually checked for each channel, and detection thresholds were adjusted. The translational status of each mRNA over time was then evaluated. Only events that remained for at least 1 min were classified as translation-ON state. To quantify translation in each reading frame, the fraction of time during which each frame was detected was calculated relative to the total duration in which translation was classified as ON.

## Results

### *ATXN8OS* undergoes AUG-independent translation in all three reading frames

To examine whether translation occurs from the *ATXN8OS* gene (Fig. 1), we constructed a nano-luciferase (Nluc)-based reporter system containing the *ATXN8OS* CUG/CUA repeat together with its 3′ UTR flanking sequence (Fig. 2A), as established in previous RAN translation studies (17,21). In this reporter, an Nluc sequence lacking an AUG start codon (GGG-Nluc) and fused to a C-terminal 3×HA tag was placed downstream of the *ATXN8OS* repeat. To enable detection of translation in all three reading frames, we inserted 1, 2, or 0 nucleotides (nt) between the repeat and Nluc (+1 nt: poly L (0), +2 nt: poly Y-poly C (+1), 0 nt: poly T-poly A (+2)).

**Figure 2.**
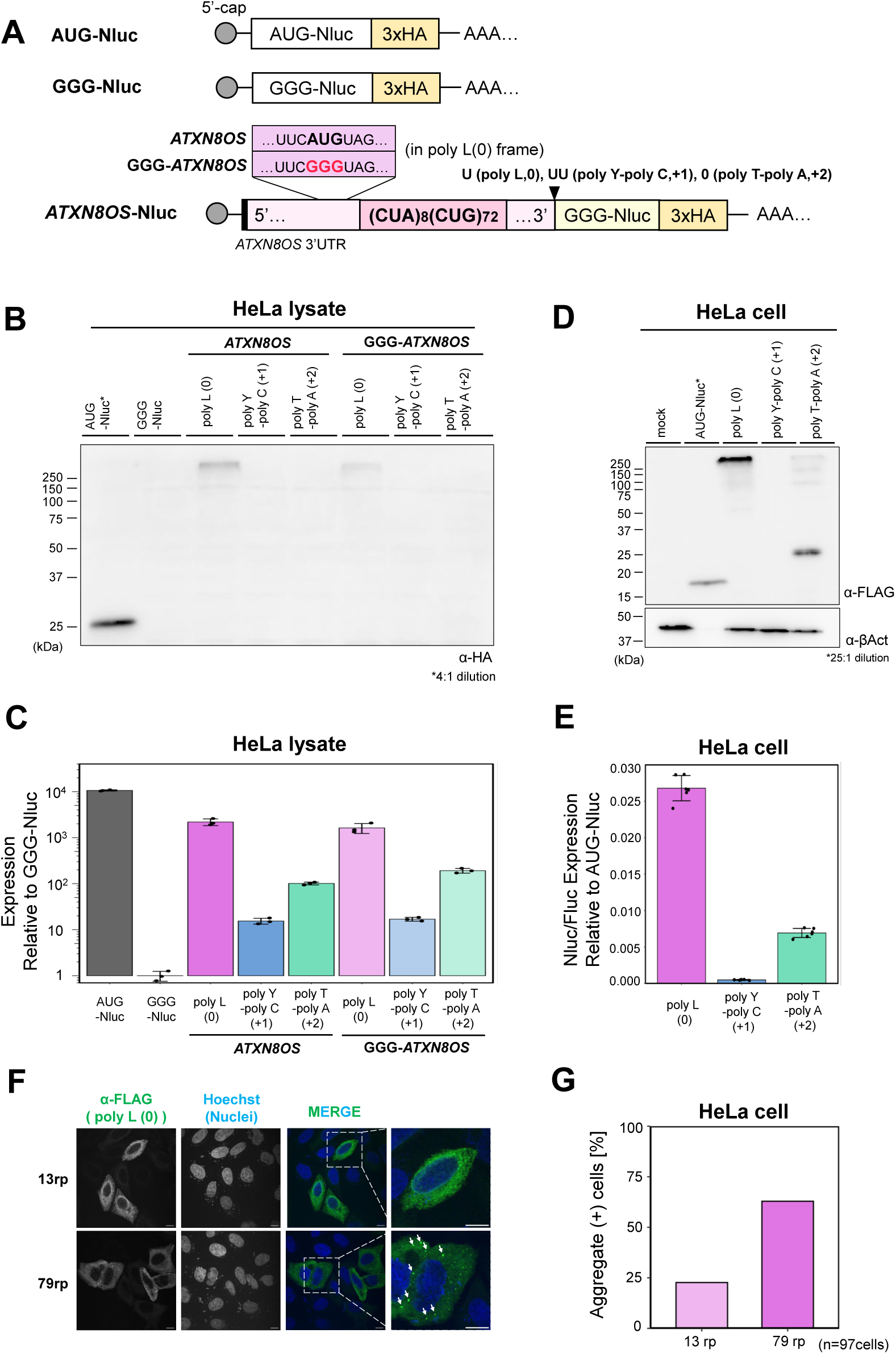
*ATXN8OS* repeat-associated translation in HeLa lysate and cells using a nano-luciferase reporter. **A:** Schematic of nano-luciferase (Nluc)-3×HA reporters. Nluc was used as a reporter enzyme to quantify translation efficiency. *ATXN8OS* and GGG-*ATXN8OS* reporters were constructed. **B:** Anti-HA Western blot of the Nluc reporter-translated HeLa lysate. Predicted molecular sizes: AUG-Nluc: 25 kDa, poly L (0): 36 kDa, poly Y-poly C (+1): 36 kDa, and poly T-poly A (+2): 34 kDa. **C:** Expression of *ATXN8OS*-Nluc reporters normalized to GGG-Nluc in HeLa lysate. Error bars represent standard deviations (±SD) from three independent experiments. **D:** Anti-FLAG Western blot of the Nluc reporter-expressed HeLa cells. Predicted molecular sizes: AUG-Nluc: 21 kDa, poly L (0): 33 kDa, poly Y-poly C (+1): 32 kDa, and poly T-poly A (+2): 30 kDa. **E:** Relative Nluc/Fluc expression of the *ATXN8OS*-Nluc reporters normalized to AUG-Nluc in HeLa cells measured by dual-luciferase assay. Error bars represent ±SD from six independent experiments. **F:** Immunostaining of *ATXN8OS*-FLAG reporter-expressing HeLa cells. Scale bar indicates 10 µm. The arrows indicate aggregates derived from poly L (0). **G:** The percentage of cells with aggregates among those showing accumulation of translation products.

We prepared capped and polyadenylated mRNAs containing the *ATXN8OS* repeat sequence, and translated them in vitro using HeLa cell lysate or RRL, followed by Western blotting analysis using the HA tag. We detected a high-molecular weight translation product from the poly L (0) frame (HeLa lysate: Fig. 2B, RRL: Supplementary Fig. S2A), suggesting that this product is aggregation-prone due to the hydrophobic nature of poly L. Although translation products from the poly Y-poly C (+1) and poly T-poly A (+2) frames were not readily detectable by Western blotting (HeLa lysate: Fig. 2B, RRL: Supplementary Fig. S2A), a luciferase reporter assay, which is more sensitive, detected expression from all three frames (HeLa lysate: Fig. 2C, RRL: Supplementary Fig. S2B). The highest translation efficiency was observed in the poly L (0) frame (∼20% in HeLa lysate and ∼40% in RRL relative to AUG-Nluc), followed by the poly T-poly A (+2) (∼1% in both lysates) and poly Y-poly C (+1) frame (∼0.1% in both lysates).

We next generated analogous reporters for expression in cultured cells and assessed their translation in HeLa and HEK293T cells. In these cellular contexts, the poly L (0) frame consistently exhibited the highest translation activity, followed by the poly T-poly A (+2) frame, whereas translation from the poly Y-poly C (+1) frame was barely detectable (HeLa: Fig. 2D, E, HEK293T: Supplementary Fig. S3A-C). These cellular results closely mirrored the trends observed in the in vitro translation assays.

We next examined the aggregation properties of poly L (0) by immunofluorescence. HeLa cells were transiently transfected with an *ATXN8OS* reporter containing 13 or 79 CUA/CUG repeat, followed by a 3×FLAG tag. Compared with the 13-repeat construct, cells expressing the 79-repeat construct displayed a markedly increased number of bright cytoplasmic foci (Fig. 2F), consistent with the aggregation-prone nature of poly L (0). Quantitatively, among cells successfully expressing the reporter, 68% of 79-repeat cells exhibited FLAG-positive foci, whereas 25% of 13-repeat cells did so (Fig. 2G). These results indicate that *ATXN8OS* translation products, including poly L, exhibit repeat length-dependent aggregation.

The *ATXN8OS* transcript contains an endogenous AUG codon located 21 nt upstream of the repeat in the poly L (0) frame (Supplementary Fig. S1B). We therefore examined whether translation could occur in the absence of this AUG codon, as reported for *ATXN8* (4). To this end, we generated a mutant reporter series (GGG-*ATXN8OS*), in which the AUG was mutated to GGG (Fig. 2A). Western blotting of the GGG-*ATXN8OS* reporters yielded results similar to those obtained with the original *ATXN8OS* reporter: poly L (0) remained readily detectable, whereas products from other frames were faint (Fig. 2B, Supplementary Fig. S2A). Luciferase assays further revealed that translation from the GGG-*ATXN8OS* reporters remained robust across all three frames (∼70% for poly L (0), ∼110% for poly Y-poly C (+1), and ∼190% for poly T-poly A (+2) relative to the original *ATXN8OS* reporter) (Fig. 2C, Supplementary Fig. S2B). Collectively, these results demonstrate that *ATXN8OS* translation occurs independently of an AUG codon, providing direct evidence that the *ATXN8OS* repeat undergoes RAN translation.

### Poly L (0) translation initiates from multiple upstream codons

We next investigated the mechanism of translation initiation in *ATXN8OS*. First, we examined whether *ATXN8OS* translation requires a 5′ m⁷G cap using HeLa lysate. Translation efficiency dropped to ∼5% in all three frames when the reporter lacked a 5′ m⁷G cap (Supplementary Fig. S4A). Moreover, the addition of excess free m⁷GpppG, which competitively inhibits eIF4E binding, strongly suppressed translation across all frames (Supplementary Fig. S4B). Together, these results demonstrate that *ATXN8OS* translation is initiated via a cap-dependent scanning mechanism.

Based on these observations, and by analogy to other RAN translation systems such as *C9orf72*, *FMR1*, and *Nop56* (15–17), we hypothesized that translation initiation occurs at a near-cognate codon upstream of the repeat via scanning (15–17). We therefore focused on the poly L (0) frame, which exhibited the highest translation efficiency. Using the GGG-*ATXN8OS* reporter to identify RAN translation initiation sites, we introduced GGG substitutions into four candidate near-cognate codons upstream of the repeat (ACG, CUG, UUG, CUG; mut1-mut4) (Fig. 3A). Western blotting and luciferase assays revealed that mut1-3 had little effect, whereas mut4 reduced translation to <10% of the unmodified GGG-*ATXN8OS* reporter (Fig. 3B, C), identifying the CUG codon at the mut4 position as the primary initiation site for poly L (0) RAN translation.

**Figure 3.**
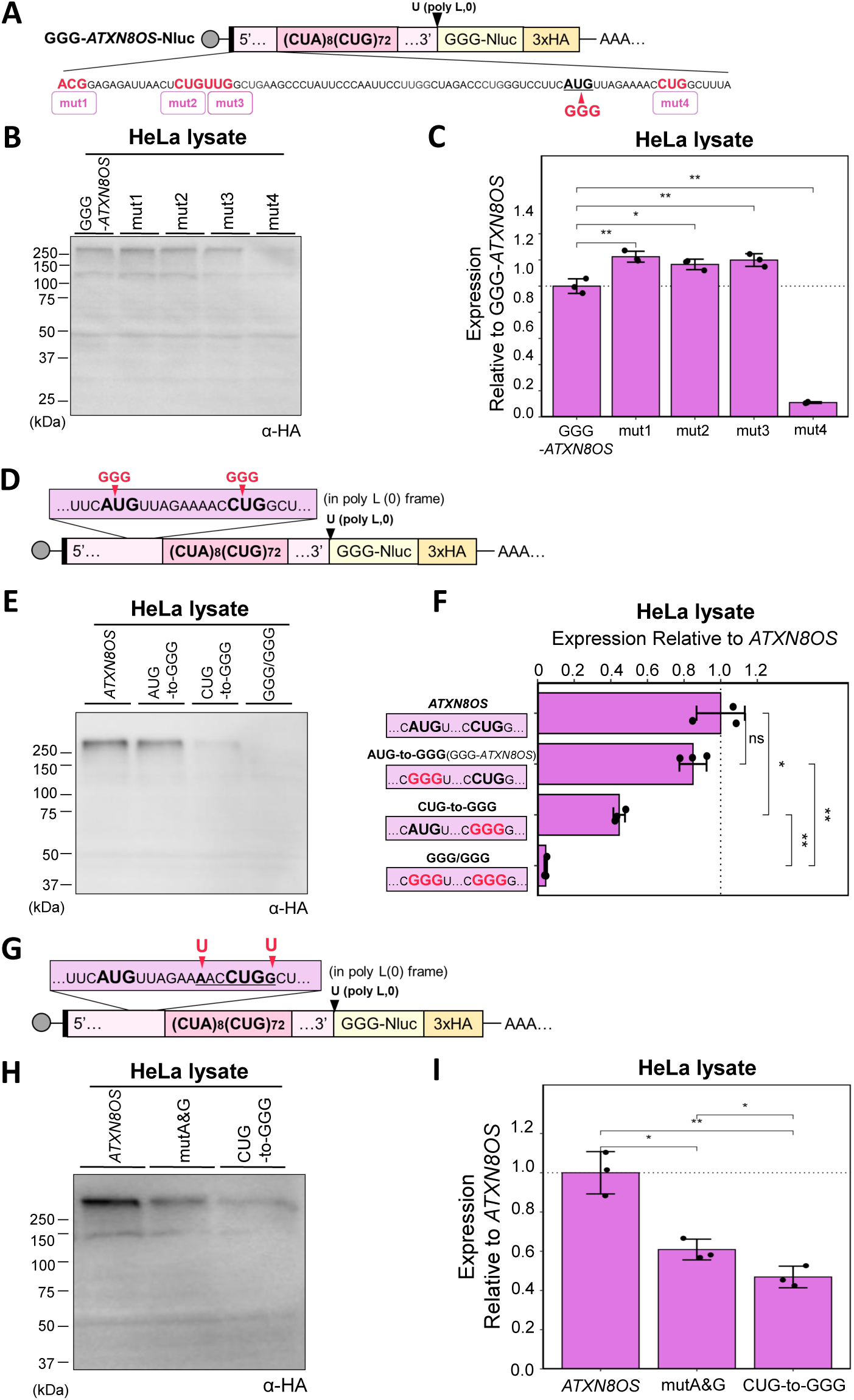
Identification of initiation sites for poly L (0) frame translation. **A, D, G:** Mutations used to identify (A, D) *ATXN8OS* poly L (0) RAN translation initiation site and (G) the effect of the Kozak sequence. **B, E, H:** Anti-HA Western blot of the Nluc reporter translated in HeLa lysate to examine (B, E) poly L (0) RAN translation initiation site and (H) the effect of the Kozak sequence. **C, F, I:** Expression of the mutant reporters relative to the expression of (C) GGG-*ATXN8OS* reporter and (F, I) *ATXN8OS* reporter in HeLa lysate. Error bars represent ±SD from three independent experiments. ∗p < 0.05; ∗∗p < 0.01; ∗∗∗p < 0.001, two-tailed Student’s *t*-test.

Mutation of the endogenous AUG codon alone partially reduced poly L (0) translation relative to the original *ATXN8OS* reporter (HeLa lysate: ∼70%; Fig. 2C, HEK293T cell: ∼50%; Supplementary Fig. S3E). Taken together, these findings indicate that both the AUG (canonical translation) and the upstream CUG codon (RAN translation) contribute to *ATXN8OS* translation initiation. To further verify this, we generated reporters carrying either single, AUG-to-GGG (=GGG-ATXN8OS) and CUG-to-GGG), or double (GGG/GGG) mutations (Fig. 3D).

Western blotting and luciferase assays showed that the CUG mutation alone reduced translation to <50% of the original *ATXN8OS* reporter, whereas the double mutation reduced translation to <10% (Fig. 3E, F). These data demonstrate that *ATXN8OS* translation is driven by both canonical AUG-initiated translation and RAN translation initiated at the upstream CUG codon. Similar results were observed in HEK293T cells (Supplementary Fig. S3D, E).

The sequence surrounding the CUG codon closely resembles a Kozak consensus motif (CC(A/G)CC<u>AUG</u>G), which is critical for translation initiation in eukaryotes (Fig. 3G, underlined) (51,52). To test whether this Kozak-like context contributes to initiation at the CUG codon, we introduced point mutations at the −3 (A-to-U) and +1 (G-to-U) positions, two key positions in the Kozak motif (mutA & G; Fig. 3G) (51,52). Western blotting and luciferase assays revealed that the mutA&G construct reduced expression to a level comparable to that of the CUG-to-GGG mutant (Fig. 3H, I). Consistent results were obtained with Kozak AUG-Nluc and Kozak CUG-Nluc reporters (Supplementary Fig. S5). These results indicate that the Kozak-like context is a critical determinant for initiation at the CUG codon in the poly L (0) frame.

### *ATXN8OS* translation is enhanced in a repeat length-dependent manner

Previous studies of RAN translation have shown that translation efficiency increases with repeat length (4,16,17,21). To test whether this property also applies to *ATXN8OS*, we designed *ATXN8OS*-Nluc reporters containing 0, 13, or 80 repeats (Fig. 4A). Luciferase assays revealed that translation in the poly L (0) frame was increased by approximately 8-fold in the 80-repeat compared with the 0-repeat construct (Fig. 4B). The poly Y-poly C (+1) and poly T-poly A (+2) frames also showed modest increases (∼1.5 fold) (Fig. 4C, D). Collectively, these results demonstrate that *ATXN8OS* translation is modulated by repeat length, consistent with a defining feature of RAN translation.

**Figure 4.**
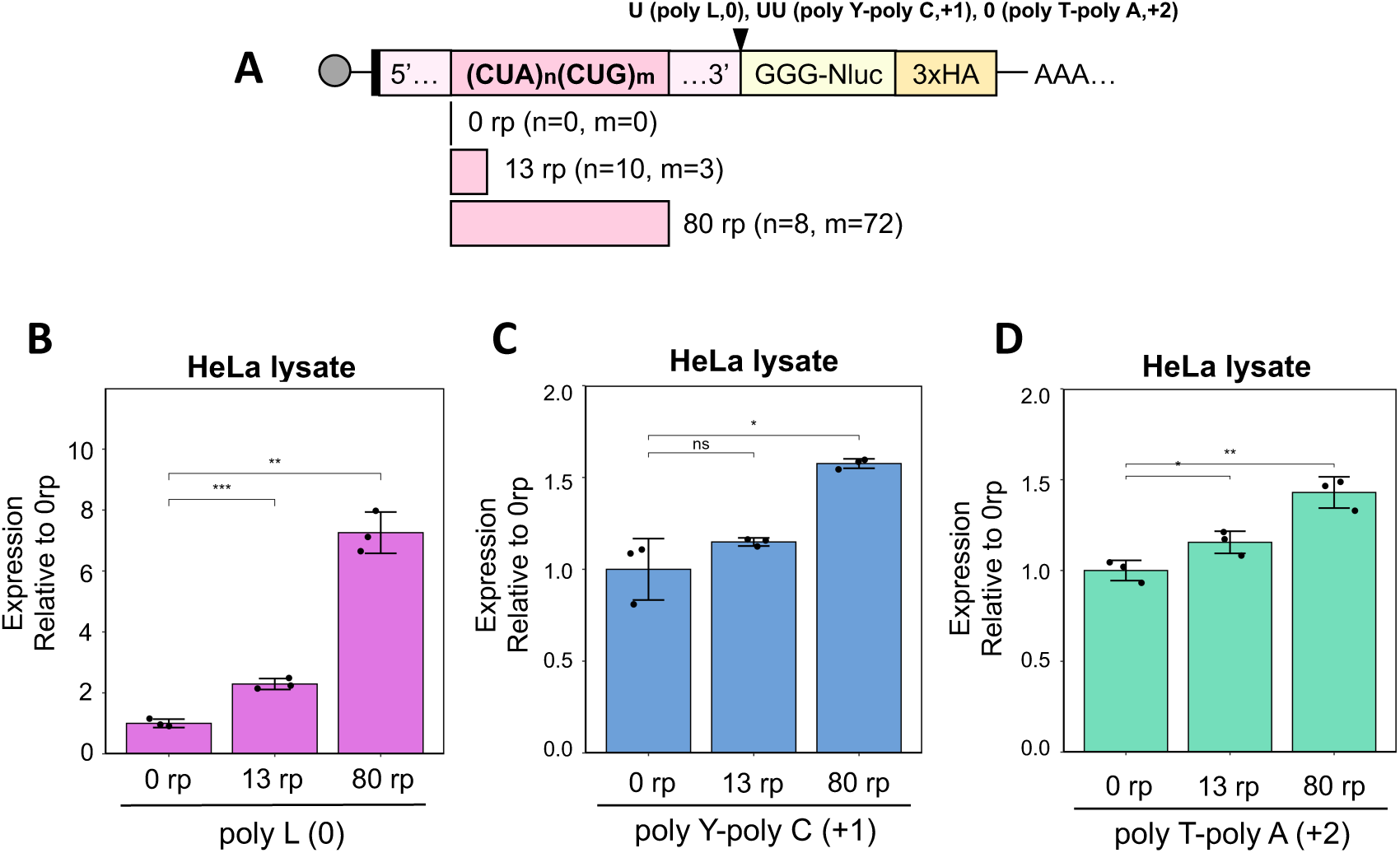
Repeat length-dependent enhancement of *ATXN8OS* translation. **A:** Schematic of the *ATXN8OS*-Nluc reporter with varying repeat numbers in each frame. **B, C, D:** Relative expression levels of *ATXN8OS*-Nluc reporters normalized to the 0-repeat (0 rp) in HeLa lysate. (B) poly L (0), (C) poly Y-poly C (+1), (D) poly T-poly A (+2) frame. Error bars represent ±SD from three independent experiments. ∗∗∗p < 0.001, one-way ANOVA with Dunnett’s multiple comparison test.

### Ribosomal frameshifting from the poly L (0) to the poly T-poly A (+2) frame occurs during *ATXN8OS* translation

A previous study reported that −1 ribosomal frameshifting (−1FS) occurs at a frequency of ∼2% at CUG 5 repeat in HEK293T cells (26). Repeat-induced ribosomal frameshifting has also been reported in other RAN translation systems, including a poly GR (+2) to poly GA (0) frameshift in C9-RAN, and a poly R (0) to poly G (+1) frameshift in FMR1-RAN (22,23). Based on these observations, we considered the possibility that ribosomal frameshifting might also occur during *ATXN8OS* translation. To test this, we used our previously established reporter system (17), in which an AUG codon was inserted upstream of the CUA/CUG repeat in frame with the poly L (0) frame (Fig. 5A). Nluc was positioned downstream of the repeat, with the insertion of 1, 2, or 0 nucleotides, enabling detection of frameshift products (+1 nt: poly L (0) to in-frame poly L (0) [No-FS], +2 nt: poly L (0) to poly Y-poly C (+1) [+1FS], 0 nt: poly L (0) to poly T-poly A (+2) [−1FS]). Translation efficiency was assessed in U2OS cells. Luciferase assays showed that, in the 80-repeat CUA/CUG construct, −1FS products were detected at ∼6.7% relative to the No-FS products, whereas in the 0-repeat construct they were detected at ∼1.9% (Fig. 5B). By contrast, +1FS products were barely detectable, consistent with a previous study (26). We also examined the potential contribution of a putative frameshifting-prone slippery sequence (T TTT TTA) immediately downstream of the CUA/CUG repeat (Supplementary Fig. S1B, underlined) and found that this sequence had no detectable effect on −1FS (Supplementary Fig. S6A, B). Together, these results demonstrate that −1 ribosomal frameshifting occurs during *ATXN8OS* translation in a repeat length-dependent manner.

**Figure 5.**
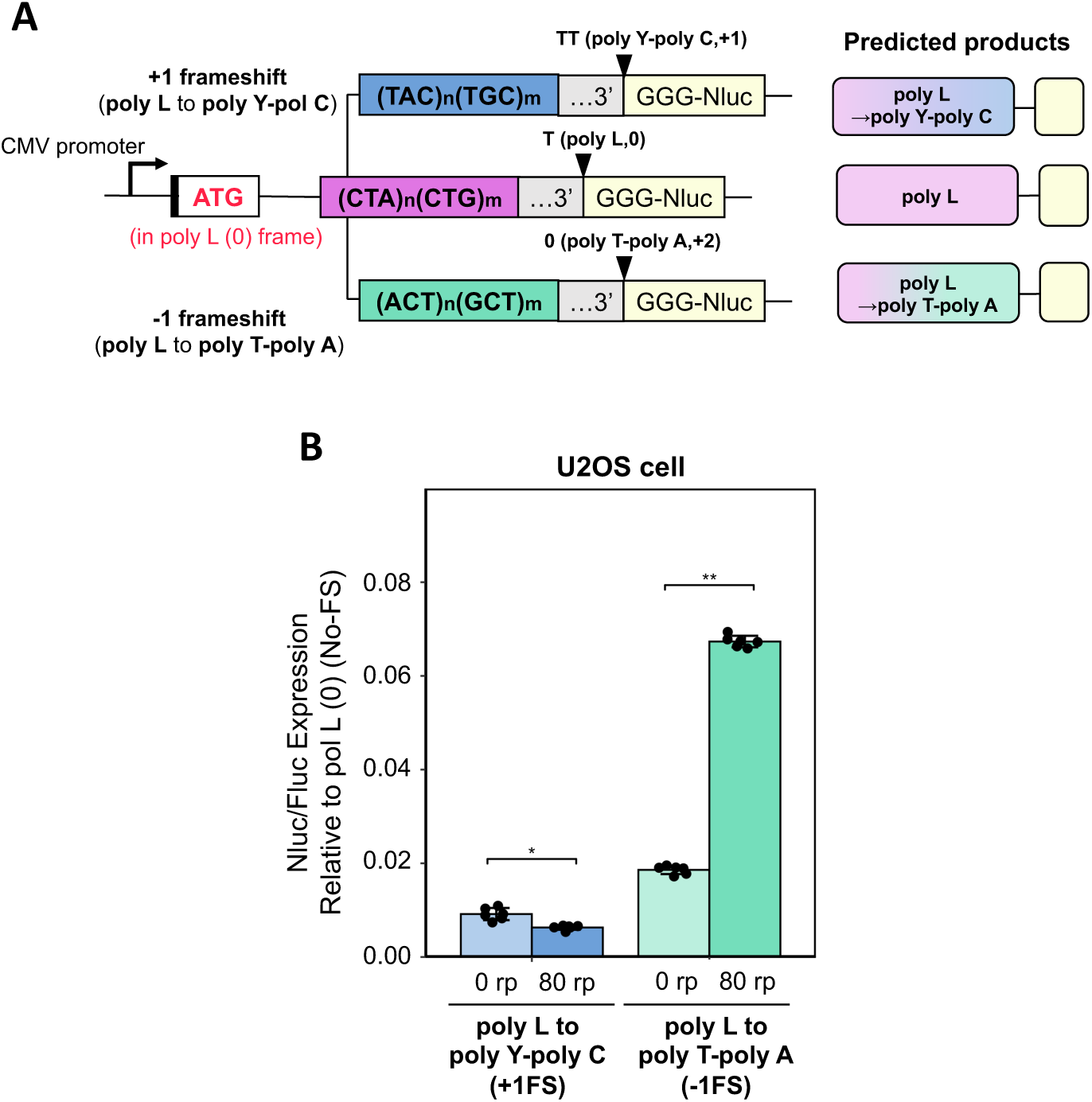
Frameshift analysis of *ATXN8OS* translation. **A:** Schematic of frameshift reporters and predicted products. AUG was inserted in poly L (0) frame, driving expression through C-terminal Nluc fused to *ATXN8OS* 3′ UTR with 0 or 80 rp. **B:** Relative Nluc/Fluc expression of frameshifted products (poly L (0) to poly Y-poly C [+1FS], poly L to poly T-poly A [−1FS]) expressed in U2OS cells. Nluc/Fluc activity for in-frame products (poly L (0) [No-FS]) is set to 1. Error bars represent ±SD from six independent experiments. ∗p < 0.05; ∗∗p < 0.01, two-tailed Student’s *t*-test.

### Live-cell imaging directly visualizes −1 frameshifting at the single-mRNA level

To directly demonstrate frameshifting in *ATXN8OS*-RAN, we applied live-cell nascent-chain tracking to visualize frameshifting events at a single-mRNA level (22,38,39). We designed a dual-color translation imaging reporter, in which the CUA/CUG repeat was inserted between a 10×spaghetti-monster HA tag (smHA) and a 36×SunTag (Fig. 6A). The poly L (0) frame was initiated from an AUG codon placed immediately upstream of smHA, while the 36×SunTag was placed in the poly T–poly A (+2) frame, followed by a stop codon. Within the 3′ UTR, 24×PP7 binding sites were inserted to visualize individual mRNAs and to tether translation sites to the plasma membrane via a CaaX-containing PP7 coat protein (PCP-CaaX (53)). We also generated an “all-probes” plasmid to label each nascent polypeptide chain and mRNA: anti-HA frankenbody–mBaoJin for detecting poly L (0) translation, anti-SunTag scFv–mScarlet for detecting poly T–poly A (+2) translation, and PCP–Halo–CaaX for labeling PP7-tagged mRNA (Supplementary Fig. S7A). These two plasmids were co-transfected into U2OS cells. Depending on the reading frame(s) used during translation, each polysome appeared as green only (poly L (0) translation / no frameshifting), magenta only (poly T–poly A (+2) translation), or a combination of both colors (frameshifting events) (Supplementary Fig. S7B).

**Figure 6.**
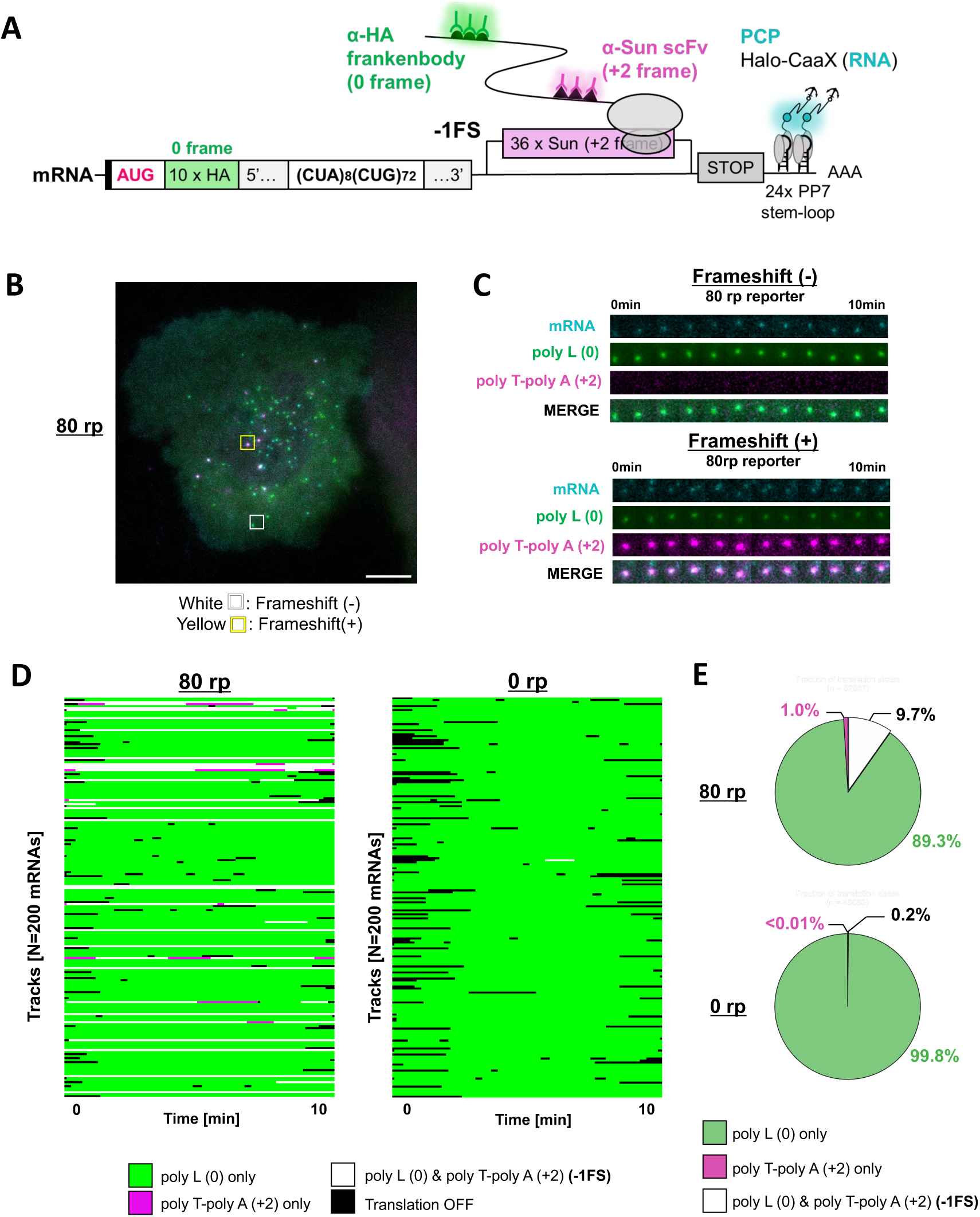
Live-cell imaging directly visualizes −1 frameshifting at the single-mRNA level. **A:** Schematic of reporters with an AUG in poly L (0) frame driving expression of an N-terminal 10×HA tag (green) in poly L (0), followed by a C-terminal 36×Sun tag (magenta) in poly T-poly A (+2) fused to the *ATXN8OS* 3’ UTR with 0 or 80 repeats. **B:** Representative image of cells expressing the 80-repeat reporter. The scale bar indicates 10 µm. Some RNA signals colocalize only with poly L (0) signals, indicating normal translation (white box). Some RNA signals colocalize with both poly L (0) and poly T-poly A (+2) signals, indicating frameshifting (yellow box). RNAs, magenta; poly L (0), blue; poly T-poly A (+2), green. For improved visibility of the image data, the contrast was adjusted. See also Supplementary Movie 3. **C:** Time-lapse images illustrating translation with or without frameshifting events. Frameshift (−) corresponds to the white box in B, and Frameshift (+) to the yellow box in B. For improved visibility of the image data, the contrast was adjusted. See also Supplementary Movie 4 and 5. **D:** Combined translating RNA tracks in 80-repeat reporter. The top 200 mRNAs ranked by tracking time are shown. (See Supplementary Fig. S7G for tracks for all mRNAs). Green, poly L (0) only; magenta, poly T-poly A (+2) only; white, both frames (−1 frameshift); black, translation off. **E:** Percentage of translating mRNAs undergoing normal translation (poly L (0) only, green), frameshifting (both frames, white), or RAN translation (poly T-poly A (+2) only, magenta).

We at first verified that internal initiation within the CUA/CUG repeat region is negligible using a negative control reporter, in which a stop codon was inserted immediately downstream of the smHA tag (80 rp-STOP), thereby eliminating any potential internal initiation (Supplementary Fig. S7C, D; Supplementary Movie 1, 2).

We then visualized the dual-color *ATXN8OS* RAN reporter in live cells. In a representative cell, we observed ∼50 mRNA spots (cyan), a subset of which colocalized with poly L (0) nascent peptide spots (green), indicating active AUG-initiated poly L (0) frame translation (Fig. 6B, white box; Fig. 6C, Frameshift (−); Supplementary Movie 3, white circle, Movie 4). Among these sites, we further identified several signals that also colocalized with poly T-poly A (+2) spots (green), revealing frameshifting events (Fig. 6B, yellow box; Fig. 6C, Frameshift (+); Supplementary Movie 3-5). Notably, some mRNAs switched from a poly L (0) translating state to a −1 frameshifting state during the observation period (Supplementary Fig. S7G, Left; Supplementary Movie 3, yellow diamond). In contrast, in the 0-repeat reporter, although mRNA spots colocalized with poly L (0) spots, no robust frameshifting sites were detected (Supplementary Fig. S7E, F; Supplementary Movies 6, 7).

We next quantified translation states over time for all translating mRNAs in our dataset. Each mRNA track was derived from quantitative analysis of signal intensities in each channel (Supplementary Fig. S7G, left). Analysis of 1,118 mRNAs in the 80-repeat reporter showed that 89.3% were in the normal AUG-initiated poly L (0) frame translation state, whereas 9.7% were in the frameshift state (Fig. 6D, E, Supplementary Fig. S7G; 80 rp). In contrast, analysis of 1,012 mRNAs in the 0-repeat reporter demonstrated that 99.8% were in the AUG-initiated poly L (0) state, while only 0.2% were in the frameshifting state (Fig. 6D, E, Supplementary Fig. S7G; 0 rp). Together with the reporter-assay results (Fig. 5B), these single-molecule observations provide direct evidence that −1 frameshifting during *ATXN8OS* translation occurs in a repeat length-dependent manner.

### The integrated stress response enhances *ATXN8OS*-RAN translation

Finally, we examined whether *ATXN8OS* translation is modulated by the integrated stress response (ISR), as reported for other RAN translation systems (11,15,17,28). To induce endoplasmic reticulum (ER) stress, we treated cells with thapsigargin (TG) (Fig. 7A). Upon TG treatment, translation in the poly L (0) and poly T-poly A (+2) frames increased by approximately 1.5-fold (Fig. 7B). These results demonstrate that *ATXN8OS* RAN translation is upregulated by ISR, consistent with previous observations in other RAN translation systems.

**Figure 7.**
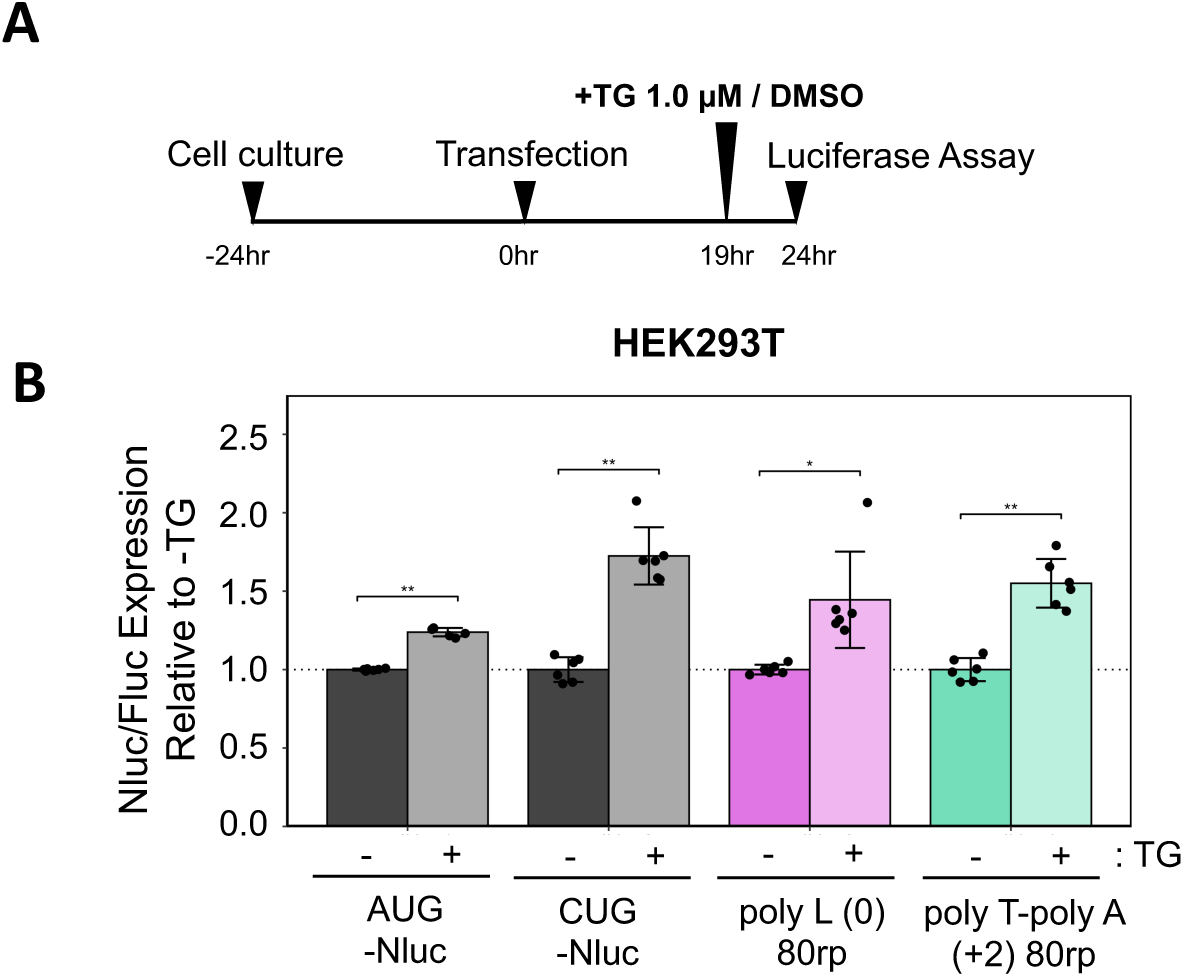
The integrated stress response enhances *ATXN8OS* translation. **A:** Schematic of stress induction by thapsigargin (TG). **B:** Relative Nluc/Fluc expression of the *ATXN8OS*-Nluc reporters normalized to HEK293T cells treated with DMSO (−TG). Error bars represent ±SD from six independent experiments. n.s., not significant; ∗∗*p* < 0.01; ∗∗∗*p* < 0.001, two-tailed Student’s *t*-test.

## Discussion

In this study, we employed a multilayered experimental strategy spanning cell-free translation systems to live-cell imaging to investigate the molecular mechanisms underlying translation of patient-derived *ATXN8OS* transcript. We establish that RAN translation occurs from the *ATXN8OS* CUA/CUG repeat within its native surrounding sequence context and dissect the underlying mechanisms. Using reporter assays, we observed that translation proceeds in all three reading frames in the absence of an AUG codon, with the poly L (0) frame showing the highest efficiency, followed by the poly T-poly A (+2) and poly Y-poly C (+1) frames (Fig. 2A-E, Supplementary Fig. S2, S3A, B, C). Mutational analyses in a cell-free translation system further identified that translation in the poly L (0) frame is initiated from both an AUG and a CUG codon (Fig. 3A-F, Supplementary Fig. S3D,E), indicating that this frame is translated via a combination of canonical AUG-initiated translation and CUG-initiated RAN translation.

Further mutational analyses demonstrated that the Kozak-like context surrounding the CUG codon is a critical determinant of its efficient use as an initiation site (Fig. 3G-I). Given that the AUG codon resides in a suboptimal Kozak context (Supplementary Fig. S1B), a downstream CUG codon embedded in a stronger Kozak context may achieve initiation efficiencies comparable to those of the AUG codon. The ability of a non-AUG codon to function as an efficient start site in the presence of an AUG codon is consistent with previous reports of alternative initiation in specific isoforms, including FGF2 and c-Myc (54–58). *ATXN8OS* translation therefore represents an additional example of non-canonical initiation that deviates from the classical rules of start codon selection. Consistent with findings from RAN translation in other genes (4,16,17,21), we observed a repeat length-dependent increase in translation efficiency. Previous studies have shown that downstream RNA secondary structures can promote translation initiation at upstream non-AUG codons, and that initiation efficiency is influenced by the distance between the start site and the structured RNA element (59,60). Notably, CUG repeats of ≥17 units are known to form more stable hairpins (61,62). Thus, the increased stability of RNA secondary structure formed by longer repeats is likely to contribute to the enhanced translation efficiency observed in our assays.

During the elongation phase, reporter assays combined with live-cell imaging revealed −1 ribosomal frameshifting from the poly L (0) to the poly T-poly A (+2) frame in a repeat length-dependent manner (Figs. 5, 6). Because this frameshifting was observed only in the presence of the repeat sequence (Supplementary Fig. S6A, B), our data suggest that, particularly in long CUG repeat RNAs (secondary structure and repetitive codon), the primary frameshift site resides within the repeat region itself. It has long been proposed that ribosomal pausing induced by higher-order RNA structures formed by repeat sequences promotes frameshifting (63,64). In addition, previous studies have shown that frameshifting can occur within continuous short CUG repeat tracts (26), indicating that repetitive codon composition alone is sufficient to stimulate ribosomal frameshifting. Together, our results agree with these findings and further suggest that, in the context of *ATXN8OS*, both the continuity of repeat codons and the formation of stable RNA secondary structures contribute to efficient −1 ribosomal frameshifting.

Live-cell imaging of frameshifting in repeat translation has been pioneered in other RAN translation systems (22,23). Our work extends this conceptual and technical framework to *ATXN8OS* to demonstrate the frameshifting on individual mRNAs. Live-cell imaging at the single-mRNA level revealed that −1 ribosomal frameshifting occurs at a frequency of ∼10% (Fig. 6E), whereas translation of the poly T–poly A (+2) product reached ∼20–30% of the poly L (0) frame level in cultured cells (Fig. 2E, Supplementary Fig. S3B). Thus, frameshifting alone cannot fully account for the poly T-poly A (+2) frame translation, indicating that additional initiation mechanisms must also contribute. Although we did not focus on the +1 frameshift from the poly L (0) to the poly Y-poly C (+1) frame, our reporter assay detected a ∼1% frameshift efficiency at 80 repeats (Fig. 5B), matching the ∼1% translation level observed in cell-based systems (Fig. 2E, Supplementary Fig. S3B). These observations suggest that the poly Y-poly C (+1) frame is predominantly generated by frameshifting events. However, it should be noted that our frameshifting assay enforces translation initiation from a canonical AUG codon and therefore does not fully recapitulate the native translational context of the *ATXN8OS* transcript.

We further explored the pathophysiological implications of *ATXN8OS* translation. Immunostaining confirmed the presence of poly L (0), which formed cytoplasmic aggregates when the repeat length was expanded to 79 repeats (Fig. 2F, G). In Huntington’s disease, RAN translation from the antisense huntingtin transcript produces poly L-containing products that are detected in caspase-3-positive brain regions, indicating the cytotoxic potential of poly L aggregates (65). By analogy, *ATXN8OS*-derived poly L (0) may similarly accumulate over time in post-mitotic neurons and thereby contribute to cellular toxicity and disease pathogenesis. Consistent with observations in other RAN translation systems (11,15,17,28), we found that *ATXN8OS* translation is upregulated under ER stress (Fig. 7). ER stress activates the integrated stress response (ISR), in which kinases such as PKR, PERK, GCN2, and HRI phosphorylate eIF2α, leading to global suppression of canonical translation (66). However, non-canonical translation pathways, including non-AUG initiation, IRES-dependent translation, and upstream ORF translation, can bypass this suppression (67–69). In agreement with this framework, both the poly L (0) and poly T-poly A (+2) translation, neither of which requires a canonical AUG codon, were similarly enhanced by ER stress. These findings raise the possibility of a feed-forward loop in which *ATXN8OS* translation products exacerbate cellular stress, thereby further activating the ISR and promoting additional RAN translation, as proposed for C9-RAN (15,18,28,70).

In summary, our results define the molecular mechanisms of *ATXN8OS* RAN translation and provide a framework for understanding how this process contributes to the pathogenesis of SCA8.

## ACKNOWLEDGEMENTS

We thank the Center for Integrative Bioscience at Science Tokyo for DNA sequencing. We thank Kodai Machida and Hiroaki Imataka for helping with the HeLa lysate, Gabriel Galindo, Hisaaki Hirose, and Hiroshi Kimura for their support and advice.

## FUNDING

This work was supported by MEXT Grants-in-Aid for Scientific Research (grant numbers JP26116002, JP18H03984, JP21H04763, and JP20H05925 to HT), AMED-CREST under Grant Number JP21gm1410008 (to H.T.), JST, the establishment of university fellowships towards the creation of science technology innovation, JSPS KAKENHI (grant numbers JPMJFS2112 and 24KJ1067 to H. Ito), Uehara Memorial Foundation (to H.T.), Mitsubishi Foundation (to H.T.), and Daiichi Sankyo (to H.T.).

## AUTHOR CONTRIBUTIONS

S.S., H.I. and M.H. performed experiments; M.H. and Y.N. provided materials; S.S., H.I., T.J.S., and H.T. conceived the study, designed experiments, analyzed the results, approved the manuscript, and are accountable for all aspects of the work; H.T. supervised the entire project; and S.S., H.I. and H.T. wrote the manuscript with the help of other authors.

## CONFLICT OF INTEREST

The authors declare that they have no conflict of interest with the contents of this article.

## DATA AVAILABILITY

Data in this manuscript have been uploaded to the Mendeley Data repository (doi: 10.17632/fkc5z63r5y.1).

## Notes

### Competing Interest Statement

The authors have declared no competing interest.

## REFERENCES

1. Malik, I., Kelley, C. P., Wang, E. T. & Todd, P. K. Molecular mechanisms underlying nucleotide repeat expansion disorders. Nat. Rev. Mol. Cell Biol. 22, 589–607 (2021).

2. Rodriguez, C. M. & Todd, P. K. New pathologic mechanisms in nucleotide repeat expansion disorders. Neurobiol. Dis. 130, (2019).

3. Todd, P. K. & Paulson, H. L. RNA-mediated neurodegeneration in repeat expansion disorders. Ann. Neurol. 67, 291–300 (2010).

4. Zu, T. et al. Non-ATG-initiated translation directed by microsatellite expansions. Proc. Natl. Acad. Sci. U. S. A. 108, 260–265 (2011).

5. Bañez-Coronel, M. et al. A pathogenic mechanism in Huntington’s disease involves small CAG-repeated RNAs with neurotoxic activity. PLoS Genet. 8, (2012).

6. Ash, P. E. A. et al. Unconventional Translation of C9ORF72 GGGGCC Expansion Generates Insoluble Polypeptides Specific to c9FTD/ALS. Neuron 77, 639–646 (2013).

7. Todd, P. K. et al. CGG repeat-associated translation mediates neurodegeneration in fragile X tremor ataxia syndrome. Neuron 78, 440–455 (2013).

8. Banez-Coronel, M. & Ranum, L. P. W. Repeat-associated non-AUG (RAN) translation: insights from pathology. Laboratory Investigation 99, 929–942 (2019).

9. Paulson, H. Repeat expansion diseases. Handb. Clin. Neurol. 147, 105–123 (2018).

10. Green, K. M., Linsalata, A. E. & Todd, P. K. RAN translation-What makes it run? Brain Res. 1647, 30–42 (2016).

11. Jazurek-Ciesiolka, M. et al. RAN Translation of the Expanded CAG Repeats in the SCA3 Disease Context. J. Mol. Biol. 432, (2020).

12. Zu, T. et al. RAN Translation Regulated by Muscleblind Proteins in Myotonic Dystrophy Type 2. Neuron 95, 1292–1305.e5 (2017).

13. Soragni, E. et al. Repeat-Associated Non-ATG (RAN) Translation in Fuchs’ Endothelial Corneal Dystrophy. Invest. Ophthalmol. Vis. Sci. 59, 1888–1896 (2018).

14. Todd, T. W. et al. Hexanucleotide Repeat Expansions in c9FTD/ALS and SCA36 Confer Selective Patterns of Neurodegeneration In Vivo. Cell Rep. 31, (2020).

15. Green, K. M. et al. RAN translation at C9orf72-associated repeat expansions is selectively enhanced by the integrated stress response. Nat. Commun. 8, (2017).

16. Kearse, M. G. et al. CGG Repeat-Associated Non-AUG Translation Utilizes a Cap-Dependent Scanning Mechanism of Initiation to Produce Toxic Proteins. Mol. Cell 62, 314–322 (2016).

17. Hasumi, M. et al. Dissecting the mechanism of NOP56 GGCCUG repeat-associated non-AUG translation using cell-free translation systems. Journal of Biological Chemistry 301, (2025).

18. Sonobe, Y. et al. Translation of dipeptide repeat proteins from the C9ORF72 expanded repeat is associated with cellular stress. Neurobiol. Dis. 116, 155–165 (2018).

19. Komar, A. A. & Hatzoglou, M. Cellular IRES-mediated translation: the war of ITAFs in pathophysiological states. Cell Cycle 10, 229–240 (2011).

20. Wang, S. et al. Nuclear export and translation of circular repeat-containing intronic RNA in C9ORF72-ALS/FTD. Nat. Commun. 12, (2021).

21. Ito, H. et al. Reconstitution of C9orf72 GGGGCC repeat-associated non-AUG translation with purified human translation factors. Sci. Rep. 13, (2023).

22. Latallo, M. J. et al. Single-molecule imaging reveals distinct elongation and frameshifting dynamics between frames of expanded RNA repeats in C9ORF72-ALS/FTD. Nat. Commun. 14, (2023).

23. Wright, S. E. et al. CGG repeats trigger translational frameshifts that generate aggregation-prone chimeric proteins. Nucleic Acids Res. 50, 8674–8689 (2022).

24. Tabet, R. et al. CUG initiation and frameshifting enable production of dipeptide repeat proteins from ALS/FTD C9ORF72 transcripts. Nat. Commun. 9, (2018).

25. Yu, C. H., Teulade-Fichou, M. P. & Olsthoorn, R. C. L. Stimulation of ribosomal frameshifting by RNA G-quadruplex structures. Nucleic Acids Res. 42, 1887–1892 (2014).

26. Ren, G. et al. Ribosomal frameshifting at normal codon repeats recodes functional chimeric proteins in human. Nucleic Acids Res. 52, 2463–2479 (2024).

27. Joazeiro, C. A. P. Ribosomal Stalling During Translation: Providing Substrates for Ribosome-Associated Protein Quality Control. Annu. Rev. Cell Dev. Biol. 33, 343–368 (2017).

28. Cheng, W. et al. C9ORF72 GGGGCC repeat-associated non-AUG translation is upregulated by stress through eIF2α phosphorylation. Nat. Commun. 9, (2018).

29. Koob, M. D. et al. An untranslated CTG expansion causes a novel form of spinocerebellar ataxia (SCA8). Nat. Genet. 21, 379–384 (1999).

30. Lau, K. K. et al. Spinocerebellar ataxia type 8: clinical features in a large family. Neurology 55, 207–209 (2000).

31. Gupta, A. & Jankovic, J. Spinocerebellar ataxia 8: Variable phenotype and unique pathogenesis. Parkinsonism Relat. Disord. 15, 621–626 (2009).

32. Samukawa, M. et al. PSP-Phenotype in SCA8: Case Report and Systemic Review. Cerebellum 18, 76–84 (2019).

33. Hirano, M. et al. Noncoding repeat expansions for ALS in Japan are associated with the ATXN8OS gene. Neurol. Genet. 4, (2018).

34. Nemes, J. P., Benzow, K. A. & Koob, M. D. The SCA8 transcript is an antisense RNA to a brain-specific transcript encoding a novel actin-binding protein (KLHL1). Hum. Mol. Genet. 9, 1543–1551 (2000).

35. Moseley, M. L. et al. Bidirectional expression of CUG and CAG expansion transcripts and intranuclear polyglutamine inclusions in spinocerebellar ataxia type 8. Nat. Genet. 38, 758–769 (2006).

36. Daughters, R. S. et al. RNA gain-of-function in spinocerebellar ataxia type 8. PLoS Genet. 5, (2009).

37. Mutsuddi, M., Marshall, C. M., Benzow, K. A., Koob, M. D. & Rebay, I. The Spinocerebellar Ataxia 8 noncoding RNA causes neurodegeneration and associates with Staufen in Drosophila. Current Biology 14, 302–308 (2004).

38. Lyon, K., Aguilera, L. U., Morisaki, T., Munsky, B. & Stasevich, T. J. Live-Cell Single RNA Imaging Reveals Bursts of Translational Frameshifting. Mol. Cell 75, 172–183.e9 (2019).

39. Morisaki, T. et al. Real-time quantification of single RNA translation dynamics in living cells. Science 352, 1425–1429 (2016).

40. Ito, H. et al. Canonical translation factors eIF1A and eIF5B modulate the initiation step of repeat-associated non-AUG translation. Nucleic Acids Res. 53, (2025).

41. Boersma, S. et al. Multi-Color Single-Molecule Imaging Uncovers Extensive Heterogeneity in mRNA Decoding. Cell 178, 458–472.e19 (2019).

42. Galindo, G., Fixen, G. M., Heredia, A., Morisaki, T. & Stasevich, T. J. All Probes Plasmids (APPs) for multicolor and long-term tracking of single-mRNA translation dynamics. Mol. Biol. Cell 36, (2025).

43. Zhang, H. et al. Bright and stable monomeric green fluorescent protein derived from StayGold. Nat. Methods 21, 657–665 (2024).

44. Zhao, N. et al. A genetically encoded probe for imaging nascent and mature HA-tagged proteins in vivo. Nat. Commun. 10, (2019).

45. Mikami, S., Kobayashi, T., Masutani, M., Yokoyama, S. & Imataka, H. A human cell-derived in vitro coupled transcription/translation system optimized for production of recombinant proteins. Protein Expr. Purif. 62, 190–198 (2008).

46. Fujino, Y. et al. FUS regulates RAN translation through modulating the G-quadruplex structure of GGGGCC repeat RNA in C9orf72-linked ALS/FTD. Elife 12, (2023).

47. Tokunaga, M., Imamoto, N. & Sakata-Sogawa, K. Highly inclined thin illumination enables clear single-molecule imaging in cells. Nat. Methods 5, (2007).

48. Pinkard, H. et al. Pycro-Manager: open-source software for customized and reproducible microscope control. Nat. Methods 18, 226–228 (2021).

49. Schindelin, J., et al. Fiji: an open-source platform for biological-image analysis. Nat. Methods 9, 676–682 (2012).

50. Ershov, D. et al. TrackMate 7: integrating state-of-the-art segmentation algorithms into tracking pipelines. Nat. Methods 19, 829–832 (2022).

51. Kozak, M. Point mutations define a sequence flanking the AUG initiator codon that modulates translation by eukaryotic ribosomes. Cell 44, 283–292 (1986).

52. Kozak, M. At least six nucleotides preceding the AUG initiator codon enhance translation in mammalian cells. J. Mol. Biol. 196, 947–950 (1987).

53. Yan, X., Hoek, T. A., Vale, R. D. & Tanenbaum, M. E. Dynamics of Translation of Single mRNA Molecules In Vivo. Cell 165, 976–989 (2016).

54. Arnaud, E., et al. A New 34-Kilodalton Isoform of Human Fibroblast Growth Factor 2 Is Cap Dependently Synthesized by Using a Non-AUG Start Codon and Behaves as a Survival Factor. MOLECULAR AND CELLULAR BIOLOGY vol. 19 505–514 (1999).

55. Hann, S. R., King, M. W., Bentley, D. L., Anderson, C. W. & Eisenman, R. N. A non-AUG translational initiation in c-myc exon 1 generates an N-terminally distinct protein whose synthesis is disrupted in Burkitt’s lymphomas. Cell 52, 185–195 (1988).

56. Prats, H. et al. High molecular mass forms of basic fibroblast growth factor are initiated by alternative CUG codons. Proc. Natl. Acad. Sci. U. S. A. 86, 1836–1840 (1989).

57. Hann, S. R. Regulation and function of non-AUG-initiated proto-oncogenes. Biochimie 76, 880–886 (1994).

58. Kearse, M. G. & Wilusz, J. E. Non-AUG translation: a new start for protein synthesis in eukaryotes. Genes Dev. 31, 1717–1731 (2017).

59. Kozak, M. Downstream secondary structure facilitates recognition of initiator codons by eukaryotic ribosomes. Proc. Nail. Acad. Sci. USA 87, 8301–8305 (1990).

60. Kozak, M. Circumstances and mechanisms of inhibition of translation by secondary structure in eucaryotic mRNAs. Mol. Cell. Biol. 9, 5134–5142 (1989).

61. Sobczak, K. et al. Structural diversity of triplet repeat RNAs. Journal of Biological Chemistry 285, 12755–12764 (2010).

62. Napierała, M. & Krzyzosiak, W. J. CUG repeats present in myotonin kinase RNA form metastable ‘slippery’ hairpins. Journal of Biological Chemistry 272, 31079–31085 (1997).

63. Giedroc, D. P. & Cornish, P. V. Frameshifting RNA pseudoknots: structure and mechanism. Virus Res. 139, 193–208 (2009).

64. Yu, C. H., Teulade-Fichou, M. P. & Olsthoorn, R. C. L. Stimulation of ribosomal frameshifting by RNA G-quadruplex structures. Nucleic Acids Res. 42, 1887–1892 (2014).

65. Bañez-Coronel, M. et al. RAN Translation in Huntington Disease. Neuron 88, 667–677 (2015).

66. Pakos - Zebrucka, K., et al. The integrated stress response. EMBO Rep. 17, 1374–1395 (2016).

67. Young, S. K. & Wek, R. C. Upstream Open Reading Frames Differentially Regulate Gene-specific Translation in the Integrated Stress Response. Journal of Biological Chemistry 291, 16927–16935 (2016).

68. Starck, S. R. et al. Translation from the 5’ untranslated region shapes the integrated stress response. Science 351, (2016).

69. Hinnebusch, A. G., Ivanov, I. P. & Sonenberg, N. Translational control by 5’-untranslated regions of eukaryotic mRNAs. Science 352, 1413–1416 (2016).

70. Westergard, T. et al. Repeat-associated non-AUG translation in C9orf72-ALS/FTD is driven by neuronal excitation and stress. EMBO Mol. Med. 11, (2019).

