## Supplemental Figures for "Mechanism of *ATXN8OS* CTA/CTG repeat-associated non-AUG translation revealed by approaches ranging from cell-free translation to live-cell imaging"

**Supplementary Figures S1-S7**

Supplementary Figure S1. *ATXN8OS* gene structure and sequence.

Supplementary Figure S2. *ATXN8OS* RAN translation in rabbit reticulocyte lysate (RRL) using nano-luciferase reporter.

Supplementary Figure S3. *ATXN8OS* RAN translation in cultured cells using nano-luciferase reporter.

Supplementary Figure S4. Cap-dependency of *ATXN8OS* translation in HeLa lysate.

Supplementary Figure S5. Kozak sequence-mutated canonical and non-canonical translation reporters.

Supplementary Figure S6. Frameshift analysis in *ATXN8OS* translation.

Supplementary Figure S7. Live-cell imaging to visualize the -1 frameshifting at a single mRNA level.

**Legends for supplementary movies 1-7**

Supplementary Movie 1. Live-cell imaging of translation from the *ATXN8OS* 80 rp-STOP reporter lacking frameshifting.

Supplementary Movie 2. Cropped images of a frameshift-negative mRNA from the *ATXN8OS* 80 rp-STOP reporter.

Supplementary Movie 3. A representative movie and corresponding translation intensity traces using the *ATXN8OS* 80 rp frameshift reporter.

Supplementary Movie 4. Cropped image of frameshift-negative mRNA.

Supplementary Movie 5. Cropped images of a frameshift-positive mRNA.

Supplementary Movie 6. Live-cell imaging of translation from the *ATXN8OS* 0 rp reporter lacking frameshifting.

Supplementary Movie 7. Cropped images of a frameshift-negative mRNA from the *ATXN8OS* 0 rp reporter.

**Supplementary Tables S1-S3 (in separate Excel files)**

Supplementary Table S1: Plasmid list.

Supplementary Table S2: Primer list.

Supplementary Table S3: Antibodies list.

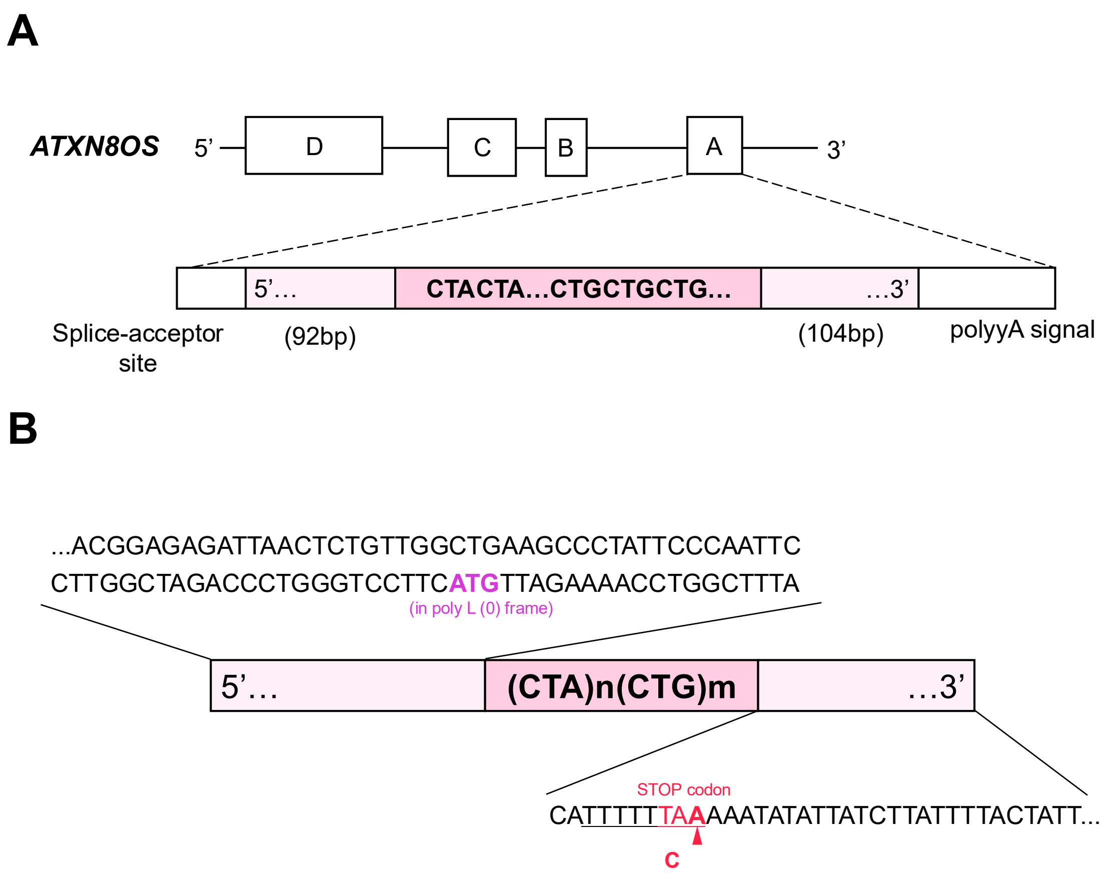


**Supplementary Figure S1. *ATXN8OS* gene structure and sequence.**

**A:** *ATXN8OS* gene structure. The CTA/CTG repeat is located in the 3′ UTR of the *ATXN8OS* gene. A, B, C, and D represent exon regions. **B:** Flanking sequence of *ATXN8OS* CTA/CTG repeat. An 85-bp upstream and a 35-bp downstream region were used for reporter constructs. An ATG codon exists upstream of the repeat sequence. The downstream TAA Stop codon was substituted with a TAC codon in the reporter. The underline indicates the slippery sequence.

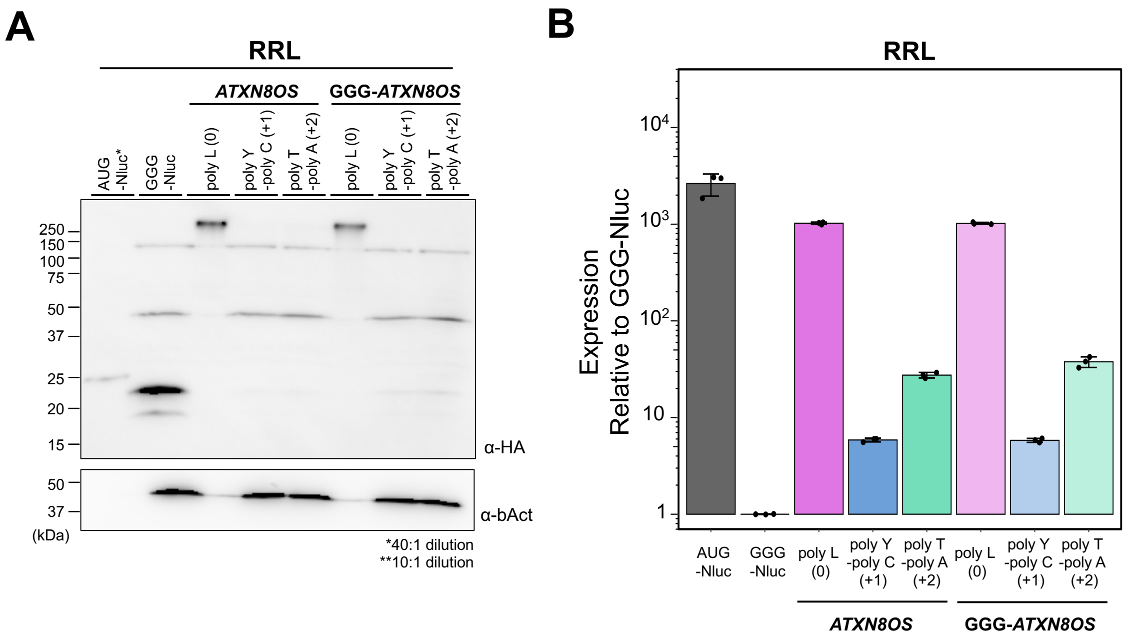


**Supplementary Figure S2.** ***ATXN8OS*** **RAN translation in rabbit reticulocyte lysate (RRL) using nano-luciferase reporters.**

**A:** Anti-HA Western blot of the Nluc reporter-translated RRL. Predicted molecular sizes: AUG-Nluc: 25 kDa, poly L (0): 36 kDa, poly Y-poly C (+1): 36 kDa, and poly T-poly A (+2): 34 kDa. Anti-β-actin (bAct) was used as a loading control. **B:** Expression of Nluc reporters normalized to GGG-Nluc in RRL. Error bars represent standard deviations (±SD) from three independent experiments.

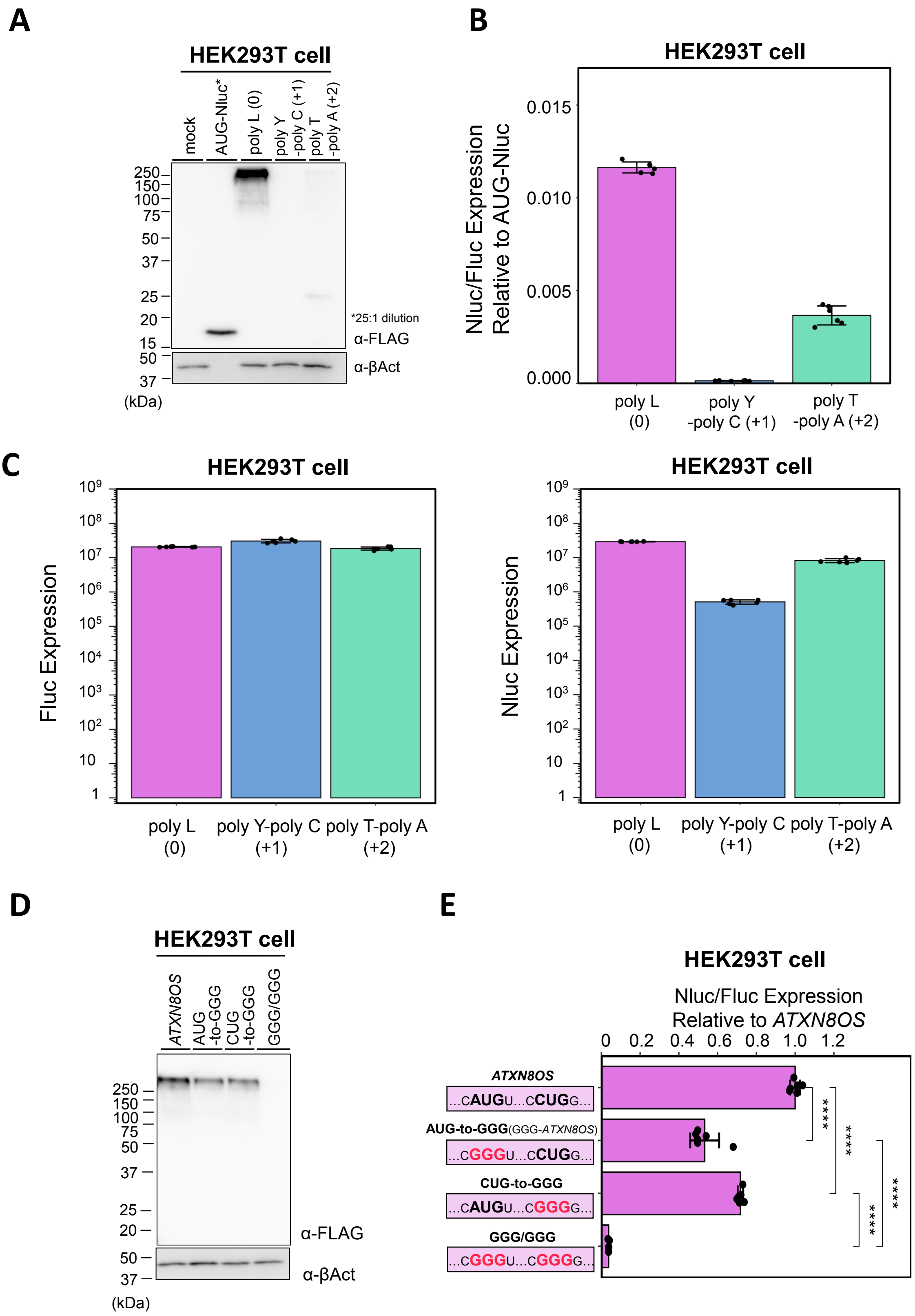
A
HEK293T cell
mock
AUG-Nluc*
poly L (0)
poly Y -poly C (+1)
poly T -poly A (+2)
250
150
100
75
50
37
25
20
15
50
37
(kDa)
*25:1 dilution
α-FLAG
α-βAct
B
HEK293T cell
Nluc/Fluc Expression Relative to AUG-Nluc
0.015
0.010
0.005
0.000
poly L (0)
poly Y -poly C (+1)
poly T -poly A (+2)
C
HEK293T cell
Fluc Expression
Nluc Expression
poly L (0)
poly Y-poly C (+1)
poly T-poly A (+2)
D
HEK293T cell
ATXN8OS
AUG -to-GGG
CUG -to-GGG
GGG/GGG
α-FLAG
α-βAct
(kDa)
E
HEK293T cell
Nluc/Fluc Expression Relative to ATXN8OS
0 0.2 0.4 0.6 0.8 1.0 1.2
ATXN8OS
...CAUGU...CCUGG...
AUG-to-GGG(GGG-ATXN8OS)
...CGGGU...CCUGG...
CUG-to-GGG
...CAUGU...CGGGG...
GGG/GGG
...CGGGU...CGGGG...
****

**Supplementary Figure S3.** ***ATXN8OS*** **RAN translation in cultured cells using nano-luciferase reporter.**

**A:** Anti-FLAG Western blot of the *ATXN8OS*-Nluc reporter expressed in HEK293T cells. Predicted molecular sizes of the products: AUG-Nluc: 21 kDa, poly L (0): 33 kDa, poly Y-poly C (+1): 32 kDa, and poly T-poly A (+2): 30 kDa. **B:** Relative expression of the ATXN8OS-Nluc reporters normalized to AUG-Nluc in HEK293T cells. Error bars represent ±SD from six independent experiments. **C:** Expression of ATG-Fluc and *ATXN8OS*-Nluc reporters plasmids expressed in HEK293T cells. ATG-Fluc control showed consistent expression across frames. For *ATXN8OS*-Nluc, poly L (0) exhibited the highest expression among the three frames, being approximately 4-fold higher than poly T-poly A (+2). Error bars represent ±SD from six independent experiments. **D:** Anti-FLAG Western blot of translation products in HEK293T cells using the candidate initiation site mutated reporters. **E:** Relative Nluc/Fluc expression of the reporters with mutations normalized to the *ATXN8OS* reporter in HEK293T cells. Error bars represent ±SD from six independent experiments. *$p < 0.05$; **$p < 0.01$; ***$p < 0.001$, two-tailed Student's t-test.

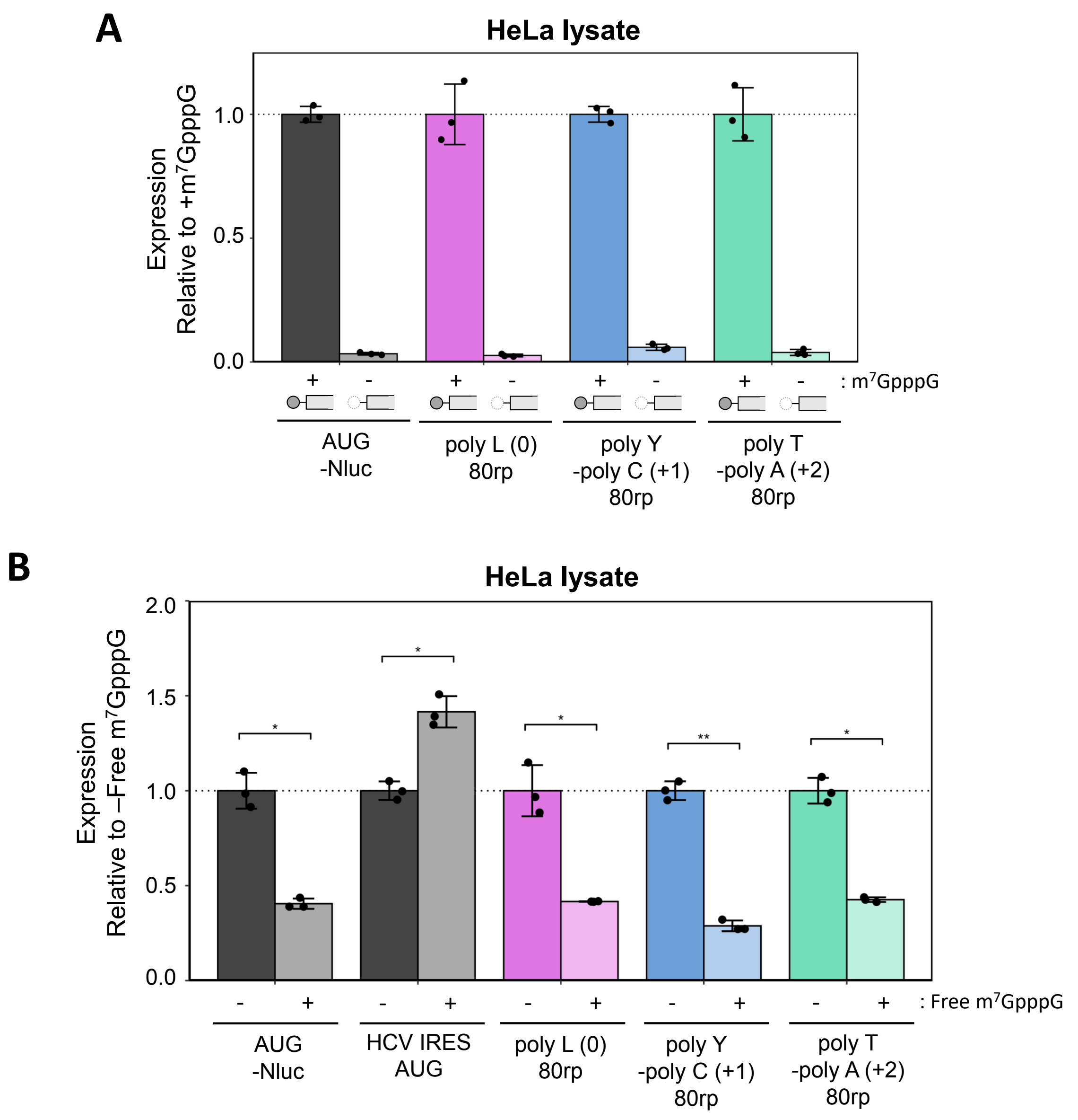


**Supplementary Figure S4. Cap-dependency of *ATXN8OS* translation in HeLa lysate.**

**A:** m7GpppG-cap dependence of *ATXN8OS*-Nluc reporter translation in HeLa lysate. Canonical AUG-initiated translation (AUG-Nluc) was included as a control.

**B:** Effect of free $m^7GpppG$ on the *ATXN8OS*-Nluc reporter translation in HeLa lysate. Nluc reporters with AUG-start and IRES were also included as controls. Error bars represent ±SD from three independent experiments. n.s. = not significant; * $p < 0.05$; ** $p < 0.001$, two-tailed Student's t-test.

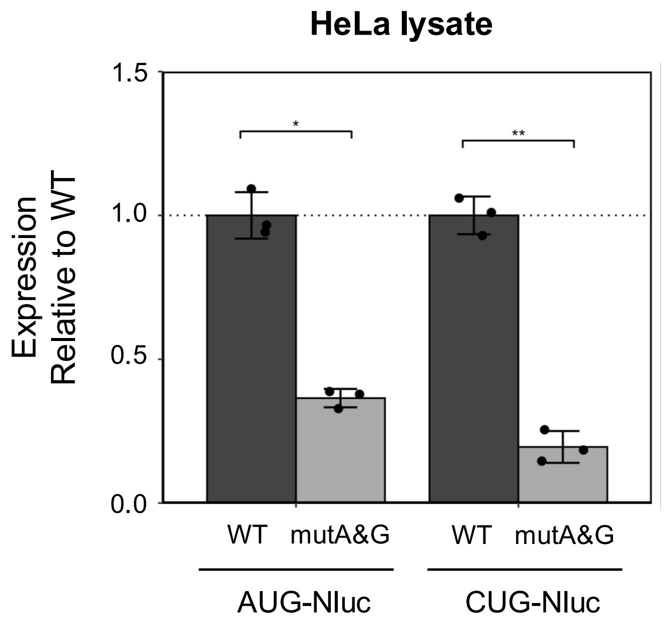


**Supplementary Figure S5. Kozak sequence-mutated canonical and non-canonical translation reporters.**

Analysis of canonical and non-canonical translation efficiencies with Kozak sequence mutation in HeLa lysate. Error bars represent ±SD from three independent experiments.

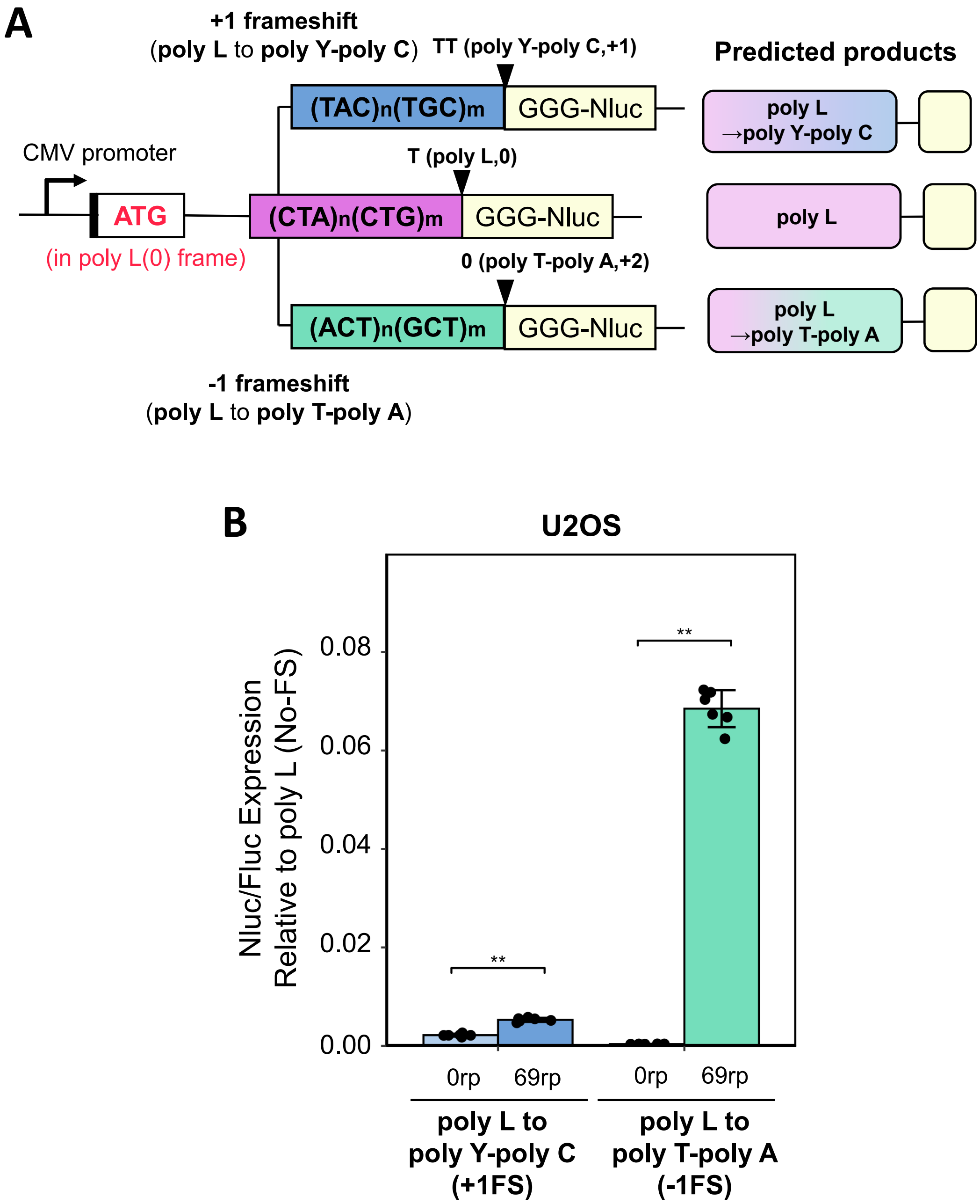


**Supplementary Figure S6. Frameshift analysis in *ATXN8OS* translation.**
**A:** Schematic of frameshift reporters and predicted products. AUG was inserted in the poly L (0) frame, driving expression through the C-terminal Nluc fused to CUA/CUG repeats (0 or 69 rp). 3′ UTR flanking sequences were all deleted. **B:** Relative Nluc/Fluc expression levels of frameshifted products (poly L (0) to poly Y-poly C (+1FS), poly L to poly T-poly A (−1FS)) expressed in U2OS cells. Nluc/Fluc activity for in-frame product (poly L (0) (No-FS)) is set to 1. Error bars represent ±SD from six independent experiments. *$p < 0.05$; **$p < 0.01$, two-tailed Student's t-test.

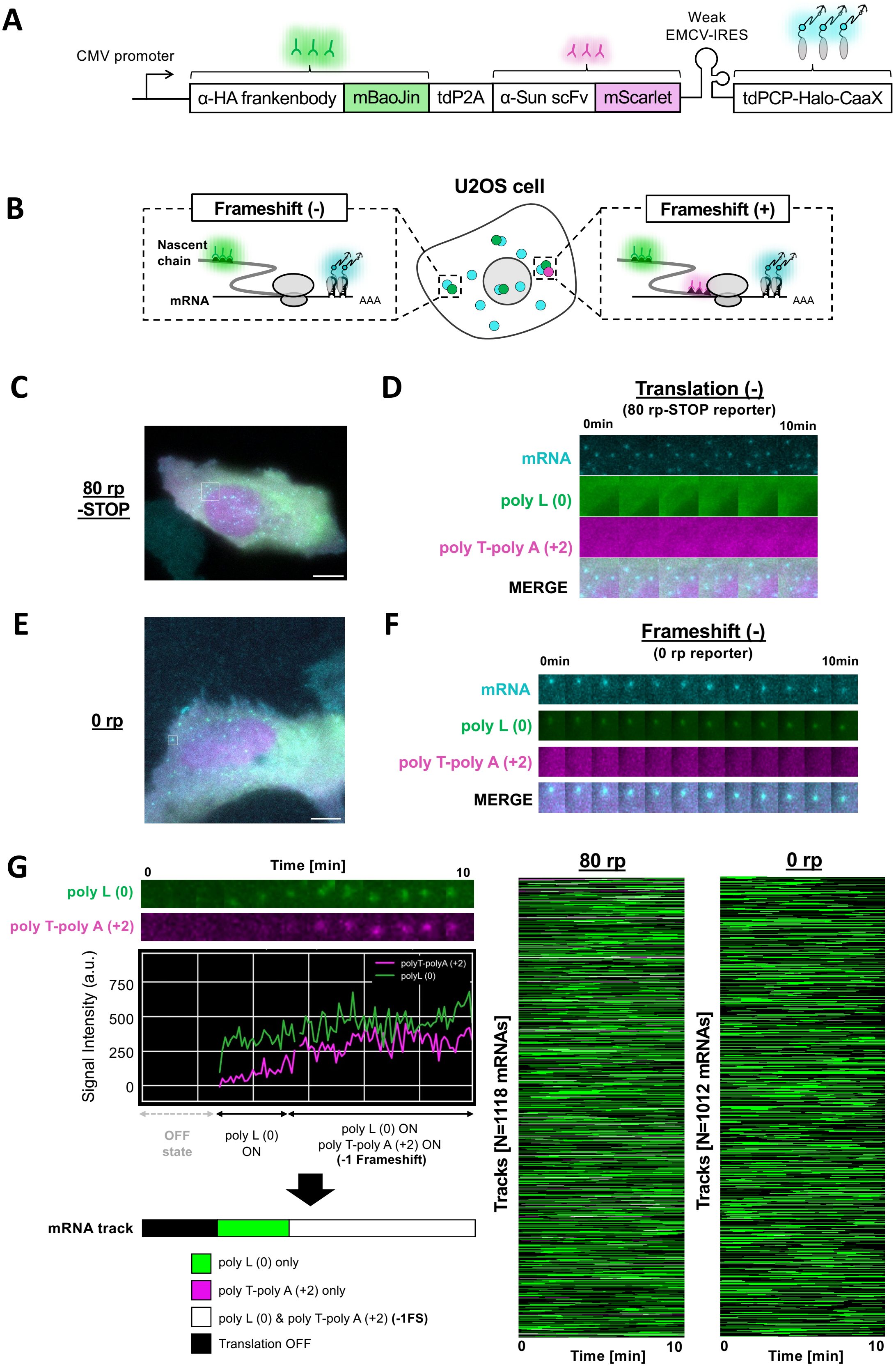

A
CMV promoter
α-HA frankenbody
mBaoJin
tdP2A
α-Sun scFv
mScarlet
Weak
EMCV-IRES
tdPCP-Halo-CaaX
B
U2OS cell
Frameshift (-)
Nascent
chain
mRNA
AAA
Frameshift (+)
AAA
C
80 rp
-STOP
D
Translation (-)
(80 rp-STOP reporter)
0min
10min
mRNA
poly L (0)
poly T-poly A (+2)
MERGE
E
0 rp
F
Frameshift (-)
(0 rp reporter)
0min
10min
mRNA
poly L (0)
poly T-poly A (+2)
MERGE
G
Time [min]
0
10
poly L (0)
poly T-poly A (+2)
Signal Intensity (a.u.)
750
500
250
0
polyT-polyA (+2)
polyL (0)
OFF
state
poly L (0)
ON
poly L (0) ON
poly T-poly A (+2) ON
(-1 Frameshift)
mRNA track
poly L (0) only
poly T-poly A (+2) only
poly L (0) & poly T-poly A (+2) (-1FS)
Translation OFF
80 rp
Tracks [N=1118 mRNAs]
0
Time [min]
10
0 rp
Tracks [N=1012 mRNAs]
0
Time [min]
10

**Supplementary Figure S7. Live-cell imaging to visualize –1 frameshifting at a single mRNA level.**

**A:** Schematic of the all-probe plasmid. By tdP2A and EMCV-IRES sequence, α-HA frankenbody-mBaojin, mScarlet and tdPCP-Halo-CaaX are expressed simultaneously. **B:** Schematic of frameshift visualization by nascent chain tracking in live cells. **C:** Representative image of 80 rp-STOP reporter-transfected cells. No RNA signals colocalized with either poly L (0) signals or poly T-poly A (+2) signals. Scale bar, 10 µm. For improved visibility of the image data, the contrast was adjusted. See also Supplementary Movie 1. **D:** Time-lapse images illustrating translation events, corresponding to the white box in C. For improved visibility of the image data, the contrast was adjusted. See also Supplementary Movie 2. **E:** Representative image of 0 rp reporter-transfected cells. Some RNA signals colocalized with poly L (0) signals, indicating translating poly L (0) translating RNA (white box). Scale bar, 10 µm. See also Supplementary Movie 6 **F:** Time-lapse images illustrating translation events, corresponding to the white box in E. For improved visibility of the image data, the contrast was adjusted. See also Supplementary Movie 7. **G:** Left: Representative analyzed mRNA track corresponding to signal intensity. Right: Combined translating RNA tracks in 80 rp (23 cells, 1,118 translating mRNAs) and 0 rp (21 cells, 1,012 translating mRNAs). Green: translating only poly L (0), Magenta: translating only poly T-poly A (+2), White: translating both frames (= –1 frameshift). Black: translation OFF.

**Legends for supplementary movies.**

**Supplementary Movie 1. Live-cell imaging of translation from the *ATXN8OS* 80 rp-STOP reporter lacking translation.**

U2OS cells were transiently transfected with the *ATXN8OS* 80 rp-STOP reporter. Cyan indicates mRNA, green indicates poly L (0) translation (smHA-Tag signal), and magenta indicates poly T–poly A (+2) translation (SunTag signal). Live-cell imaging shows no active poly L (0) translation and poly T–poly A (+2) translation. Images were acquired every 5 s, and the movie shows a 600-s time window. Scale bar, 10 µm. For improved visibility of the image data, the contrast was adjusted.

**Supplementary Movie 2. Cropped images of translation-negative mRNAs from the *ATXN8OS* 80 rp-STOP reporter.**

Cropped live-cell images of translation-negative mRNAs from the *ATXN8OS* 80 rp-STOP reporter, corresponding to the mRNA highlighted by the white box in Supplementary Fig. S7C. The movie shows a 600-s time window. Panels 1–3 show individual channels (1: mRNA; 2: poly T–poly A (+2); 3: poly L (0)), and Panel 4 shows the merged image. For improved visibility of the image data, the contrast was adjusted.

**Supplementary Movie 3. A representative movie and corresponding translation intensity traces using the *ATXN8OS* 80 rp frameshift reporter.**

U2OS cells were transiently transfected with the *ATXN8OS* frameshift reporter. Cyan: mRNA (PCP-Halo signal), green: poly L (0) (smHA-Tag signal); magenta: poly T-poly A (+2) (SunTag signal). Left: Live-cell image of a cell expressing the 80 rp frameshift reporter. A white circle represents an actively translating poly L (0) frame mRNA, and a yellow diamond represents an actively translating frameshifting (poly L (0) & poly T-poly A (+2)) mRNA. Right: Signal intensity traces of the corresponding mRNA. “Frameshift absent” corresponds to the mRNA indicated by the white circle, and “Frameshift present” corresponds to the mRNA indicated by the yellow diamond. Images were acquired every 5 s, and the movie shows a 10-min time window. Scale bar, 10 µm. For improved visibility of the image data, the contrast was adjusted.

**Supplementary Movie 4. Cropped images of a frameshift-negative mRNA.**

Cropped live-cell images of a frameshift-negative (–) mRNA corresponding to the mRNA highlighted by the white box in Fig. 6B. The movie shows a 600-s time window. Panels 1–3 show individual channels (1, mRNA; 2, poly T–poly A (+2); 3, poly L (0)), and Panel 4 shows the merged image. For improved visibility of the image data, the contrast was adjusted.

**Supplementary Movie 5. Cropped images of a frameshift-positive mRNA.**
Cropped live-cell images of frameshift-positive (+) mRNA corresponding to the mRNA highlighted by the yellow box in Fig. 6B. The movie shows a 600-s time window. Panels 1–3 show individual channels (1: mRNA; 2: poly T–poly A (+2); 3: poly L (0)), and Panel 4 shows the merged image. For improved visibility of the image data, the contrast was adjusted.

**Supplementary Movie 6. Live-cell imaging of translation from the *ATXN8OS* 0 rp reporter lacking frameshifting.**
U2OS cells were transiently transfected with the *ATXN8OS* 0 rp reporter. Cyan indicates mRNA, green indicates poly L (0) translation (smHA-Tag signal), and magenta indicates poly T–poly A (+2) translation (SunTag signal). Live-cell imaging shows active poly L (0) translation, whereas no detectable poly T–poly A (+2) translation is observed, indicating the absence of frameshifting. Images were acquired every 5 s, and the movie shows a 600-s time window. Scale bar, 10 µm. For improved visibility of the image data, the contrast was adjusted.

**Supplementary Movie 7. Cropped images of a frameshift-negative mRNA from the *ATXN8OS* 0 rp reporter.**
Cropped live-cell images of an actively translating poly L (0) mRNA from the *ATXN8OS* 0 rp reporter, corresponding to the mRNA highlighted by the white box in Supplementary Fig. S7E. The movie shows a 600-s time window. Panels 1–3 show individual channels (1: mRNA; 2: poly T–poly A (+2); 3: poly L (0)), and Panel 4 shows the merged image. For improved visibility of the image data, the contrast was adjusted.
